# Highly conserved brain vascular receptor ALPL mediates transport of engineered viral vectors across the blood-brain barrier

**DOI:** 10.1101/2024.03.12.584703

**Authors:** Tyler C. Moyer, Brett A. Hoffman, Weitong Chen, Ishan Shah, Xiao-Qin Ren, Tatiana Knox, Jiachen Liu, Wei Wang, Jiangyu Li, Hamza Khalid, Anupriya S. Kulkarni, Munachiso Egbuchulam, Joseph Clement, Alexis Bloedel, Matthew Child, Rupinderjit Kaur, Emily Rouse, Kristin Graham, Damien Maura, Zachary Thorpe, Ambreen Sayed-Zahid, Charlotte Hiu-Yan Chung, Alexander Kutchin, Amy Johnson, Johnny Yao, Jeffrey Thompson, Nilesh Pande, Mathieu E. Nonnenmacher

**Author notes:** These authors contributed equally to this work (Tyler C. Moyer, Brett A. Hoffman).

## Abstract

Delivery of systemically administered therapeutics to the central nervous system (CNS) is restricted by the blood-brain barrier (BBB). Bioengineered Adeno-Associated Virus (AAV) capsids have been shown to penetrate the BBB with great efficacy in mouse and non-human primate models, but their translational potential is often limited by species selectivity and undefined mechanisms of action. Here, we apply our RNA-guided TRACER AAV capsid evolution platform to generate VCAP-102, an AAV9 variant with markedly increased brain tropism following intravenous delivery in both rodents and primates. VCAP-102 demonstrates a similar CNS tropism in cynomolgus macaque, african green monkey, marmoset and mouse, showing 20- to 400-fold increased transgene expression across multiple brain regions relative to AAV9. We demonstrate that the enhanced CNS tropism of VCAP-102 results from direct interaction with alkaline phosphatase (ALPL), a highly conserved membrane-associated protein expressed on the brain vasculature. VCAP-102 interacts with human, primate and murine ALPL isoforms, and ectopic expression of ALPL is sufficient to initiate receptor-mediated transcytosis of VCAP-102 in an in vitro transwell model. Our work identifies VCAP-102 as a cross-species CNS gene delivery vector with a strong potential for clinical translation and establishes ALPL as a brain delivery shuttle capable of efficient BBB transport to maximize CNS delivery of biotherapeutics.

## Introduction

Gene transfer with recombinant Adeno-Associated Viruses (AAVs) holds great promise for one-time treatment of otherwise intractable inherited or idiopathic diseases, and hundreds of clinical trials have established the potential of AAVs as gene delivery vectors^1–4^. The success of Zolgensma for the treatment of infants with spinal muscular atrophy (SMA) constituted a major milestone in the establishment of AAV vectors as clinical modalities and showed that intravenously (IV) administered AAV9 capsids could cross the Blood-Brain Barrier (BBB) and transduce cells in the central nervous system (CNS)^5,6^. However, transport of natural AAV capsids across the brain vasculature is inefficient, necessitating high doses or invasive delivery for effective gene transfer, and treatment is restricted to infants up to age 2. Consequently, the gene therapy field has used AAV capsid engineering, and particularly directed evolution^7,8^, to generate mutant capsids with increased transport across the BBB. Continuous improvements in the capsid diversification and selection methods have led to the discovery of capsid variants showing dramatic improvements of BBB penetration in rodents^9–11^ or non-human primates (NHPs)^12–15^. One major limitation to the transition of these early discoveries to the clinic is the inconsistency of capsid phenotype across species, and therefore the uncertainty that capsids identified in model animals will translate in humans.

Engineered AAV capsids typically show a strong species restriction and do not recapitulate their properties between rodents and primates^9,16,17^, or even across divergent primate species such as macaques and marmosets^12,13,15^.

In parallel, recent efforts aimed at understanding the mechanism of action of CNS-trophic capsids have identified the attachment receptors used by rodent capsids to traverse the BBB via receptor-mediated transcytosis (RMT). The three vascular receptors identified so far are LY6A^18,19^, LY6C1^20,21^ and CA4^21^, used by AAV-PHP.B, 9P39 and 9P36^10^ capsids, respectively. All three receptors belong to the glycosylphosphatidylinositol (GPI)-anchored protein family and show a very low degree of conservation across species. So far, no AAV capsid variant has shown a clear cross-species translational potential and a known mechanism of action for entering the primate CNS.

Here, we describe VCAP-102, a cross-species CNS-targeting AAV capsid isolated from an AAV9 peptide display library using TRACER™ neuron-specific RNA-guided evolution^10^ in cynomolgus macaque. VCAP-102 shows high BBB penetration and CNS transduction following IV dosing in adult old-world primates (cynomolgus macaque and African Green monkey), new world primates (marmoset) and mouse. VCAP-102 transduces glial cells and neurons in all tested species with a 20- to 400-fold higher efficiency than AAV9, together with reduced liver delivery in all animal models. We identify the Tissue-Nonspecific Alkaline Phosphatase isozyme (TNAP or ALPL, OMIM 171760), a GPI-anchored protein encoded by the ALPL gene, as the vascular receptor utilized by VCAP-102 to cross the BBB. We show that ectopic expression of ALPL in a polarized cell layer is sufficient to mediate transcytosis of VCAP-102, and provide in vivo evidence that capsid-receptor interaction is directly correlated with brain transduction. Collectively, our work identifies a cross-species, neurotrophic viral vector with strong translational potential, and identifies ALPL, a highly conserved membrane-anchored receptor, as an efficient shuttle for transporting therapeutics across the BBB.

## Results

### Discovery of a cross-species BBB-penetrant AAV capsid

Towards creating an AAV peptide display library with optimal structural and functional properties, we scanned the entire surface of the AAV9 capsid to identify the most permissive locations for random peptide insertion. We used our RNA-based TRACER™ capsid engineering platform to generate an AAV9 “hotspot” library where all capsid surface positions were tested for insertion of a random 6-amino acid (AA) peptide and assigned a unique barcode (Extended Data Fig. 1a,b). The library was used to generate a mixed virus progeny and tested for transduction in various cell culture models. We observed that the VR-IV region, contained within the 3-fold axis spike of the capsid, was highly permissive and allowed random 6-mer peptide insertions at multiple locations with no discernable impact on capsid assembly or transduction (Extended Data Fig. 1c-g). For the TRACER-NHP library, random 6-AA peptides were inserted downstream of eight consecutive positions in the VR-IV loop (AA453 to 460, AAV9 VP1 numbering). After two rounds of *in vivo* selection and capsid mRNA recovery from brain tissue of adult cynomolgus macaques (*Macaca fascicularis*), we identified 1500 variants of interest and synthesized a focused library^10,22^ for selection in macaques and in C57Bl/6J and BALB/c mouse strains (Fig. 1a). The variants with the highest performance in macaque brain, designated VCAP-101 and VCAP-102, also showed the highest performance in mice (Fig. 1b,c). High-performance variants shared a common serine-proline-histidine (SPH) motif at 456-458 position, followed by a positively charged amino acid (K or R) in one of the three following amino acids (Fig. 1e), suggesting a common mechanism of BBB crossing. The 3D capsid structure of VCAP-102 was determined by cryo-electron microscopy. The capsid structure was identical to AAV9 (PDB ID 7MT0) with the exception of a protruding density on the apex of VR-IV at the 3-fold axis, consistent with the peptide insertion site (Fig. 1e). The protruding density was only visible using a 10-Å low-pass filter, suggesting a predominantly disordered state.

**Fig. 1.**
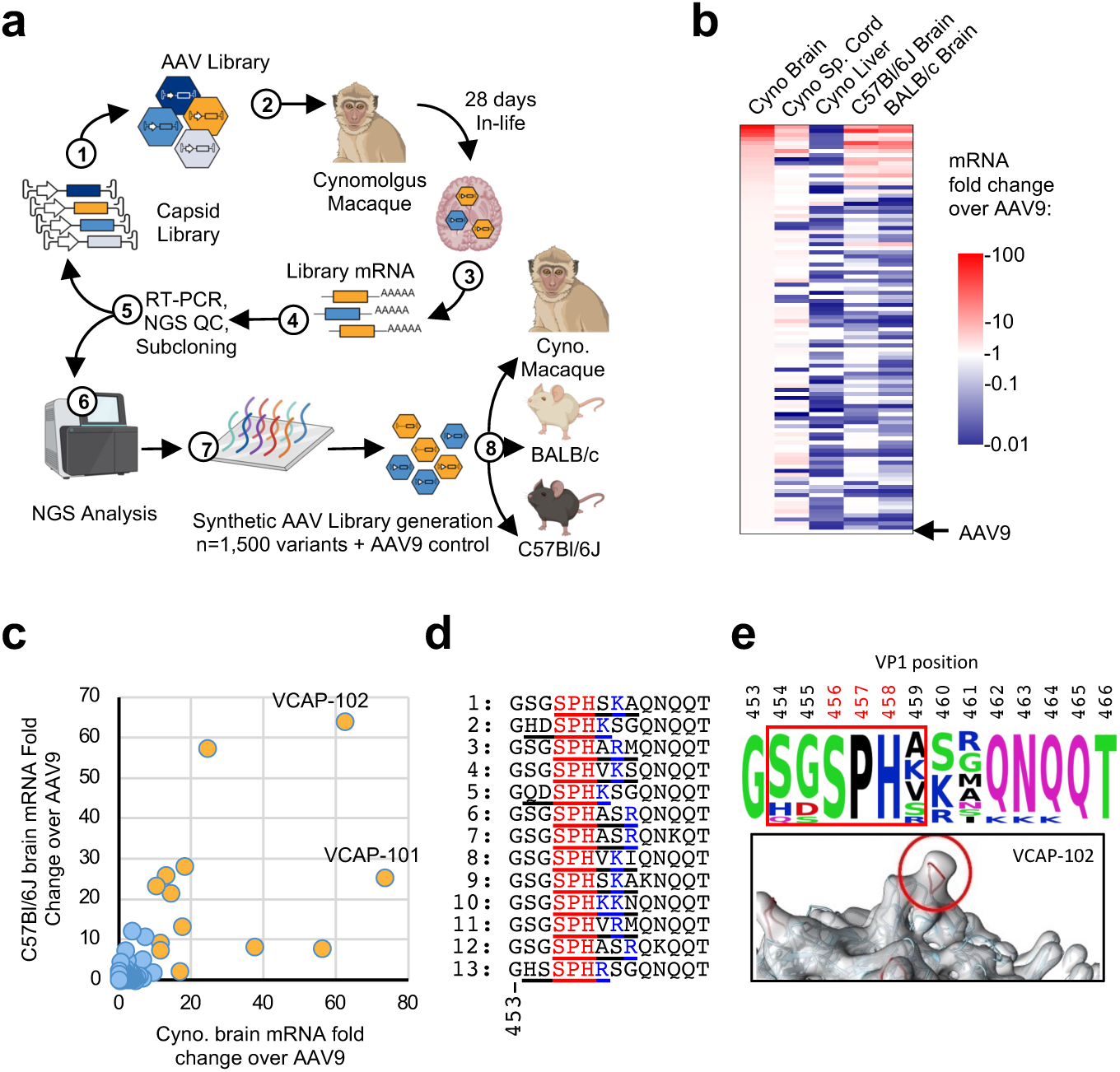
TRACER-based in vivo screen design and top hits. **a**, Design of TRACER-NHP directed evolution pipeline. **b**, Heat map of top 100 most enriched capsids in macaque brain from step (8). Color scale represents the mRNA enrichment score of the top 100 capsid versus AAV9 in indicated tissues. SC: Spinal Cord. **c**, Comparative performance of 1500 capsids in cynomolgus macaque and C57Bl/6J mouse brain. Capsids with >10-fold enrichment relative to AAV9 in macaque brain are highlighted in orange. **d**, VR-IV peptide inserts from top capsids identified in (**c**). The location of the 6mer insert is underlined, the conserved SPH motif and proximal positively charged residues are highlighted in red and blue, respectively. **e**, Position and sequence conservation of the peptides on top capsids. Upper panel: Consensus sequence of top 13 sequences from (**c**). Red positions 456-8 indicate the conserved SPH motif. Bottom panel: 3D structure of the VR-IV loop of VCAP-102 obtained by cryo-EM. The inserted 6-AA peptide is highlighted in red and corresponds to the boxed positions 454-459 in the upper panel.

### In vivo properties of VCAP-102 in rodents

To examine the performance of VCAP-101 and VCAP-102 in mouse brain we dosed BALB/c mice intravenously with each capsid containing a self-complementary (scAAV)^23,24^ ZsGreen-HA transgene under a ubiquitous promoter at a dose of 1E13 Viral Genomes per kg (VG/kg). VCAP-101 and VCAP-102 showed a broad distribution throughout the entire brain and spinal cord, outperforming AAV9 by approximately 30- and 40-fold, respectively, in terms of viral DNA biodistribution and transgene RNA expression (Fig. 2a-c). Furthermore, both variants had largely reduced expression in the liver, with VCAP-102 showing 14-fold lower gene expression than AAV9 (Fig. 2a,d,e). Transduction of the heart, skeletal muscle and kidney did not show major differences between AAV9 and VCAP-102 (Extended Data Fig. 2). Given the high CNS transduction and liver detargeting observed with VCAP-102, this capsid variant became the primary focus of our study. Next, we characterized the tropism and potency of VCAP-102 carrying a single-stranded (ssAAV) genome encoding a nuclear-localized reporter for accurate quantitation. We used a HA-tagged histone transgene (H3F3-HA) due to its strong nuclear retention and stability. We observed a dose-response across a 2-log dose range in mouse (1E12 to 1E14 VG/kg), without reaching saturation at the maximal dose (Fig. 2f,g). At the maximal tested dose VCAP-102 could transduce up to 40% of total cells in the cortex (Fig. 2g), with an even distribution between neurons and astrocytes, identified by NeuN and Sox9 markers, respectively (Fig. 2h,i; Supplementary Table 1).

**Fig. 2.**
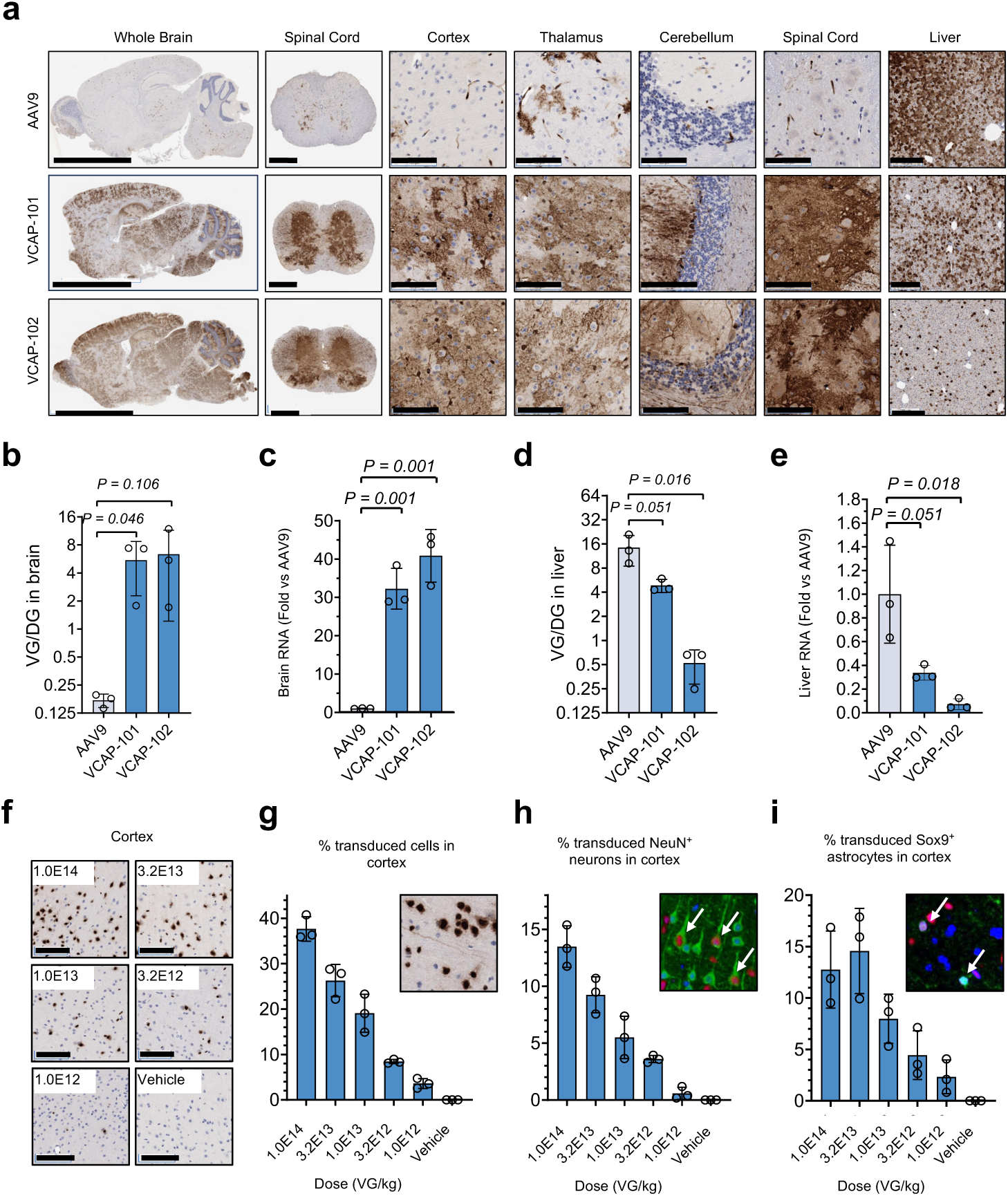
VCAP-101 and VCAP-102 display increased CNS tropism in the mouse brain. **a**, Transduction of CNS and liver in BALB/c mice. Capsids with a scAAV-CAG-ZsGreen-HA transgene were dosed IV at 1E13 VG/kg. Tissues were stained with an anti-HA antibody 28 days after IV injection. Scale bars: 5mm (whole brain), 500 µm (whole spinal cord), 100 µm (cortex, thalamus, cerebellum, spinal cord), 250 µm (liver). **b**, **c**, qPCR analysis of transgene DNA (**b**) and RNA (**c**) in the brain. DNA values represent viral genomes per diploid genome (VG/DG), RNA values are normalized to AAV9. **d**, **e**, Transgene DNA (**d**) and RNA (**e**) in the liver. **f**, Dose-dependent transduction in mouse cortex after IV injection of VCAP-102 containing a ssAAV-CAG-H3F3-HA transgene. HA tag was detected by immunohistochemistry. Scale bars: 100 µm. **g**, Percent transduced cells in the cortex from (**f**) measured by HA:hematoxylin co-localization. **h, i,** Percent transduced neurons (**h**) or astrocytes (**i**) measured by colocalization of HA tag (red) with NeuN or Sox9, respectively (green). Insets in (**g-i)** show examples of IHC and IF stainings. Arrows indicate co-labeled cells. Each data point represents an individual mouse, all plotted values represent mean ± SD (n=3). *P* values are derived from an unpaired two-tailed *t*-test.

### *In vivo* properties of VCAP-102 in primates

To test the performance of VCAP-102 capsids in primates, African Green monkeys (*Chlorocebus sabaeus*) were dosed via IV injection with 1E13 VG/kg AAV9 or VCAP-102 containing a self-complementary histone transgene (H2B-HA) under a ubiquitous CAG promoter. Broad transgene expression was observed across a range of brain regions (Fig. 3a). VCAP-102 delivered 1-2 viral genomes (VGs) per cell on average across multiple brain areas, outperforming AAV9 by 4- to 24-fold, and was capable of expressing 16- to 186-fold more transgene RNA (Fig. 3b,c). Conversely, and similar to its phenotype in rodents, VCAP-102 showed a markedly reduced liver accumulation relative to AAV9 (Fig. 3a,d,e). High-throughput analysis of immunohistochemistry stainings indicated that VCAP-102 was capable of targeting upwards of 50% of cells in the brain (Fig. 3f,g), including both astrocytes and neurons (Fig. 3h, Extended Data Fig. 3). In contrast with our mouse observations, VCAP-102 demonstrated a bias towards Sox9(+) astrocytes over neurons, labeled with either NeuN or SMI311 (Fig. 3h, Extended Data Fig. 3). Relative to AAV9, VCAP-102 showed a slight increase in heart transduction, and conversely a slightly reduced tropism for the skeletal muscle and dorsal root ganglia (Extended Data Fig. 4). VCAP-102’s performance was also examined in new world primates (marmosets, *Callithrix jacchus*) by intravenous co-injection of AAV9 and VCAP-102, each at 2E12 VG/kg, carrying a self-complementary H2B-HA or H2B-myc transgene, respectively. VCAP-102 delivered upwards of 280-fold more viral genomes and expressed 500-fold higher transgene RNA levels than AAV9 (Extended Data Fig. 5a-c). Significant liver de-targeting of VCAP-102 was also observed in this species (Extended Data Fig. 5d-f).

**Fig. 3.**
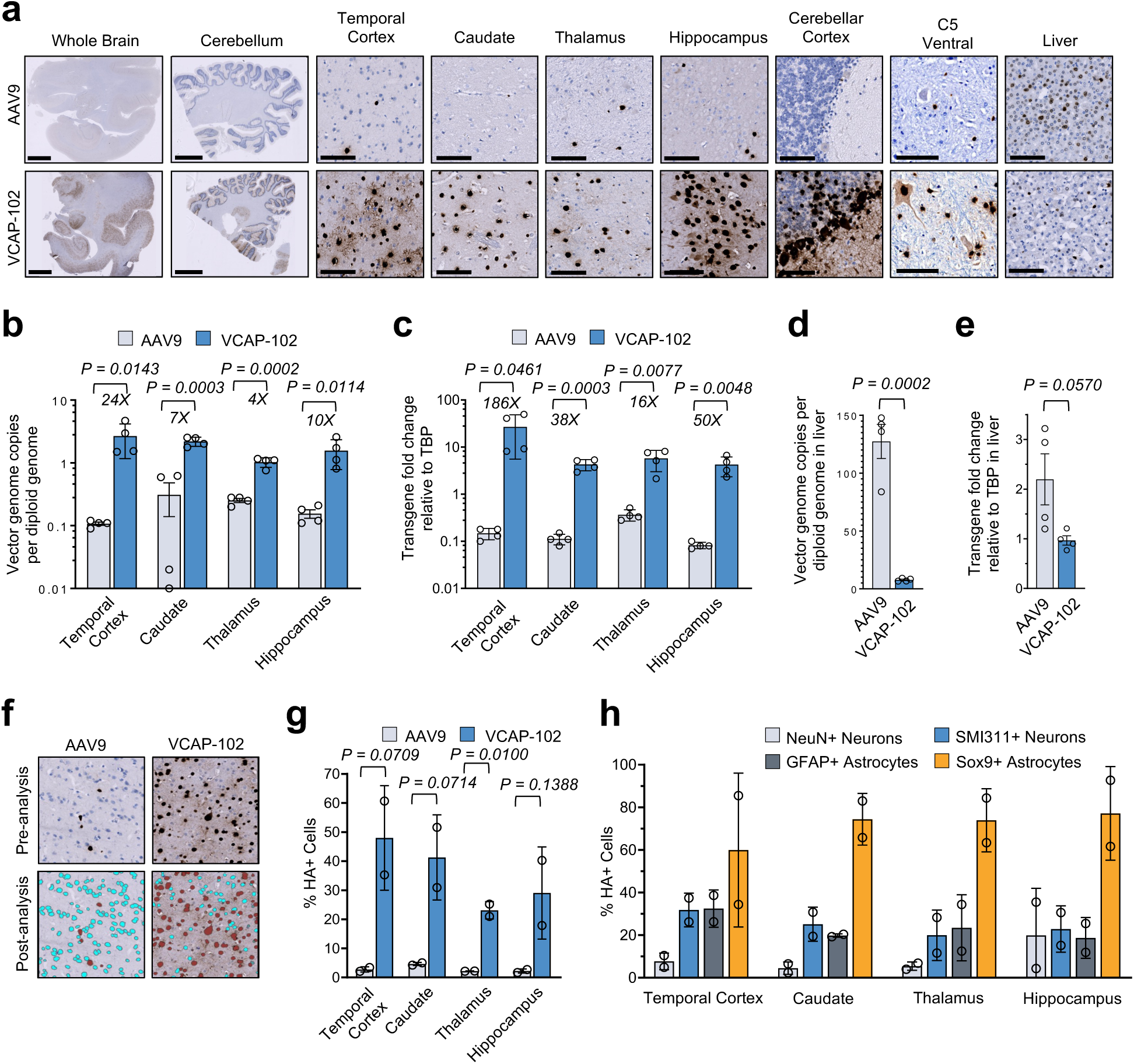
Improved CNS tropism of VCAP-102 in African green monkey. **a**, Representative images of primate CNS transduction. Images show HA tag staining in African green monkeys 28 days after IV injection of 1E13 VG/kg AAV9 or VCAP-102 capsid containing a scAAV-CAG-H2B-HA transgene. Scale bars: 5 mm for whole brain and cerebellum, 100 µm for all others. **b**, Viral DNA in indicated brain regions measured by ddPCR. Values indicate mean ± SD (n = 3) VG per cell. **c**, Real-time RT-PCR analysis of transgene mRNA expression in the brain relative to TBP housekeeping gene (mean ± SD, n = 3). **d**, VG quantification in liver. Values indicate mean ± SD (n = 3) VG per cell. **e**, Transgene mRNA expression in liver. Values indicate mean ± SD (n = 3). **f**, Automated high-throughput quantitation of transduced cells measured by co-localization of nuclear H2B-HA and hematoxylin. **g**, Quantification of HA+ nuclei in the indicated brain regions. **h**, Percentage of HA+ cells among cells positive for the indicated marker in monkeys dosed with VCAP-102. Plotted data in (**g,h**) represent one slice per monkey (n=2). Quantitative image analysis was performed on 1E3 to 1E5 cells according to region size. All *P* values are derived from an unpaired two-tailed *t*-test. Values in (**b**, **c**, **d**, **e**) represent n=4, 2 biopsy punches measured per monkey (n=2).

### Identification of ALPL as a cell surface receptor for VCAP-102

To understand the mechanism of action of VCAP-101/102 capsids and evaluate the translatability of our findings in humans, we sought to identify the vascular receptor used to transport capsids across the BBB. For this we undertook two orthogonal approaches, a capsid binding screen using Retrogenix cell microarray of human surface proteins and a transduction screen using cells stably transduced with a lentiviral human ORFeome library (Extended Data Fig. 6a,b). Both assays identified the alkaline phosphatase, biomineralization associated protein (ALPL, also known as Tissue Non-specific Alkaline Phosphatase, TNAP) encoded by the *ALPL* gene, as the most specific interaction partner. ALPL is a highly conserved GPI-anchored cell surface protein with closely related orthologs across mammals, and is expressed in multiple tissues including liver, bone, kidney and brain^25^. In the brain, ALPL expression is mainly detected in endothelial cells^26^ (Extended Data Fig. 6c), supporting a possible role in BBB transport.

To confirm ALPL functions as a primary receptor for our capsids, HEK293T cells were transfected with ALPL expression vector, followed by transduction with AAV9, VCAP-101 or VCAP-102. A plasmid encoding AAVR, a well-characterized AAV entry factor^27^, was used as a positive control. Overexpression of human ALPL in HEK239T cells led to a specific increase in VCAP-101 and VCAP-102 (Fig. 4a).

**Fig. 4.**
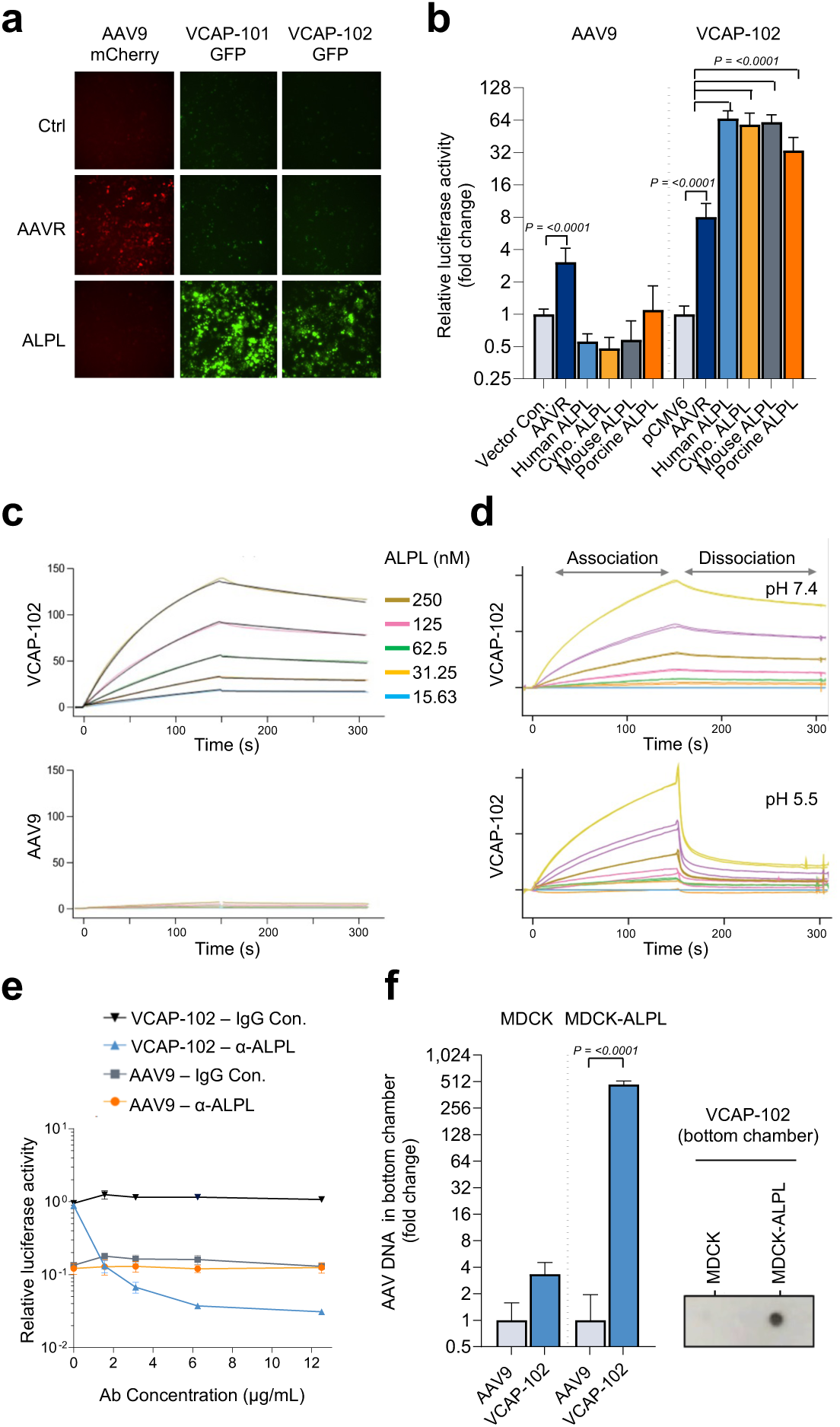
Direct ALPL:VCAP-102 interaction mediates viral transduction and transcytosis. a, Ectopic expression of ALPL increases VCAP-101 and VCAP-102 transduction. HEK293T cells transfected with indicated plasmid were treated with AAV9-mCherry or VCAP-102-GFP. Fluorescence was visualized after 24h. **b**, Functional interaction of VCAP-102 with ALPL across species. HEK293T cells transfected with indicated AAVR or ALPL plasmids were transduced with AAV9- or VCAP-102-luciferase. Data: mean ± SD luciferase activity normalized to control plasmid (n=3). **c**, Direct binding of VCAP-102 to human ALPL. Binding kinetics between VCAP-102 and human ALPL were analyzed by surface plasmon resonance (SPR). VCAP-102 or AAV9 capsids were immobilized and ALPL was used as an analyte. **d**, pH-dependent dissociation of ALPL:VCAP-102 complex, measured by SPR. VCAP-102 capsid was immobilized and ALPL was injected at pH 7.4. The dissociation was then performed at pH 7.4 or pH 5.5. **e,** Blocking of VCAP-102 transduction by ALPL antibody. HeLa cells were incubated with anti-ALPL antibody or an isotype control before adding 1E4 VG/cell AAV9 or VCAP-102 expressing luciferase. Luminescence data (mean ± SD) was normalized to untreated cells. **f**, ALPL mediates capsid transcytosis. MCDK or MDCK-ALPL cells grown on a transwell insert and AAV9 or VCAP-102 were added to the top chamber. AAV genomes in the bottom chamber was quantified 24h later by qPCR. Data indicate mean ± SD normalized to AAV9 (n=3). Insert shows detection of VCAP-102 capsid in the bottom chamber by dot blot with an anti-AAV9 antibody.

Consistent with our *in vivo* data, ALPL isoforms from human, murine and macaque origin, which share over 92% amino acid identity, showed a similar capacity to enhance of VCAP-102 transduction (Fig. 4b). We also observed functional interaction of VCAP-102 with porcine ALPL, albeit with a slightly reduced activity, suggesting that VCAP-102 could be effectively used in pig models. Conversely to our overexpression experiments, we observed that depletion of endogenous ALPL with siRNA in HeLa cells, which naturally express the receptor, significantly reduced VCAP-102 transduction (Extended Data Fig. 6d). We observed direct capsid-receptor interaction in a cell-free assay using Surface Plasmon Resonance (SPR) analysis with recombinant ALPL, showing that VCAP-102 directly interacts with ALPL with an affinity (KD) of ∼20nm at pH7.4 (Fig. 4c). Rapid dissociation of the ALPL-VCAP-102 complex was observed at pH 5.5 (Fig. 4d), which could point to a mechanism of capsid release on the basolateral side of brain endothelial cells triggered by the acidification of endosomes along the endocytosis/transcytosis pathway. The role of ALPL as a primary cell surface attachment receptor for VCAP-102 was confirmed by preincubation of HeLa cells with an anti-ALPL antibody, which dramatically reduced VCAP-102 transduction while it had no effect on AAV9 (Fig. 4e). The importance of ALPL localization to the cell surface was further confirmed by enzymatic removal of membrane-associated GPI-AP with the GPI-cleaving enzyme phosphatidylinositol-specific phospholipase C (PI-PLC)^28^, and by overexpression of a natural ALPL isoform deficient in cell membrane localization (Extended Data Fig. 6e,f). In both experiments, selective impairment of surface-localized ALPL had a negative impact on VCAP-102 transduction.

We established a capsid transcytosis model using Madin-Darby Canine Kidney cells (MDCK) in a transwell migration assay. Ectopic expression of human ALPL induced a dramatic >100-fold increase in the transcytosis of VCAP-102 while it had no measurable impact on AAV9 migration (Fig. 4f). Overall, our observations suggest that VCAP-102 binding to ALPL can trigger internalization, transcytosis and release of capsids along the endosomal trafficking pathway, in a similar fashion to BBB carriers harnessing the transferrin receptor for CNS transport^29–32^. Importantly, we observed that VCAP-102 still requires AAVR for effective transduction (Extended Data Fig. 7a) and can utilize galactose for cell attachment (Extended Data Fig. 7b), similar to its parent capsid AAV9^33,34^. This indicates that ALPL functions as an independent attachment factor for VCAP-102 while preserving the receptor usage of AAV9.

### Impact of ALPL on VCAP-102 CNS transduction *in vivo*

We sought to determine whether variation of ALPL expression level would impact VCAP-102 tropism *in vivo*. ALPL expression on the brain vasculature has been shown to increase with age in mice^26^. To test whether this age-dependent increase in ALPL expression would result in a corresponding increase in VCAP-102 transduction, young (6-8 week) and aged (∼18 month) C57Bl/6J mice were dosed intravenously with 1E13 VG/kg of AAV9 or VCAP-102. Relative to young animals, aged mice displayed a 1.5-fold higher ALPL mRNA expression (Fig. 5a) and a 1.8-fold increase in viral transgene expression throughout the entire brain and spinal cord (Fig. 5b,c), supporting a direct correlation between ALPL expression level and CNS tropism of VCAP-102 *in vivo*. We also took advantage of available small-molecule inhibitors of ALPL to investigate the impact of ALPL blocking *in vivo*. Incubation of HeLa cells with SBI-425, a small molecule inhibitor of ALPL^35^, significantly inhibited VCAP-102 transduction (Fig. 5d). The mechanism of SBI-425 is not fully elucidated, but our *in silico* modeling using DiffDock suggests that SBI-425 blocks ALPL-capsid interaction by competitive binding to the same receptor pocket (Extended Data Fig. 8a). Consistent with this hypothesis, we found that SBI-425 efficiently blocked the cell attachment and transcytosis of VCAP-102 (Extended Data Fig. 8b,c). Given the capacity of ALPL small-molecule inhibitors to block VCAP-102 attachment in cultured cells, we investigated the impact of ALPL inhibition *in vivo* by dosing VCAP-102 to mice pre-treated with SBI-425. The 9P36 capsid^10^ was chosen as a control since it is known to use CA4 as a BBB receptor and should be unaffected by the ALPL inhibitor^21^. Pre-treatment with SBI-425 resulted in a 2.5-fold reduction in brain luminescence in mice dosed with VCAP-102-luciferase at 7 days post AAV administration, whereas the 9P36-dosed mice were unaffected (Fig. 5e). At 14 days post AAV administration, SBI-425 caused a 4.1-fold reduction in VCAP-102 transgene expression in the CNS (Fig. 5f), further supporting the central role of ALPL in VCAP-102 BBB transport and CNS transduction.

**Fig. 5.**
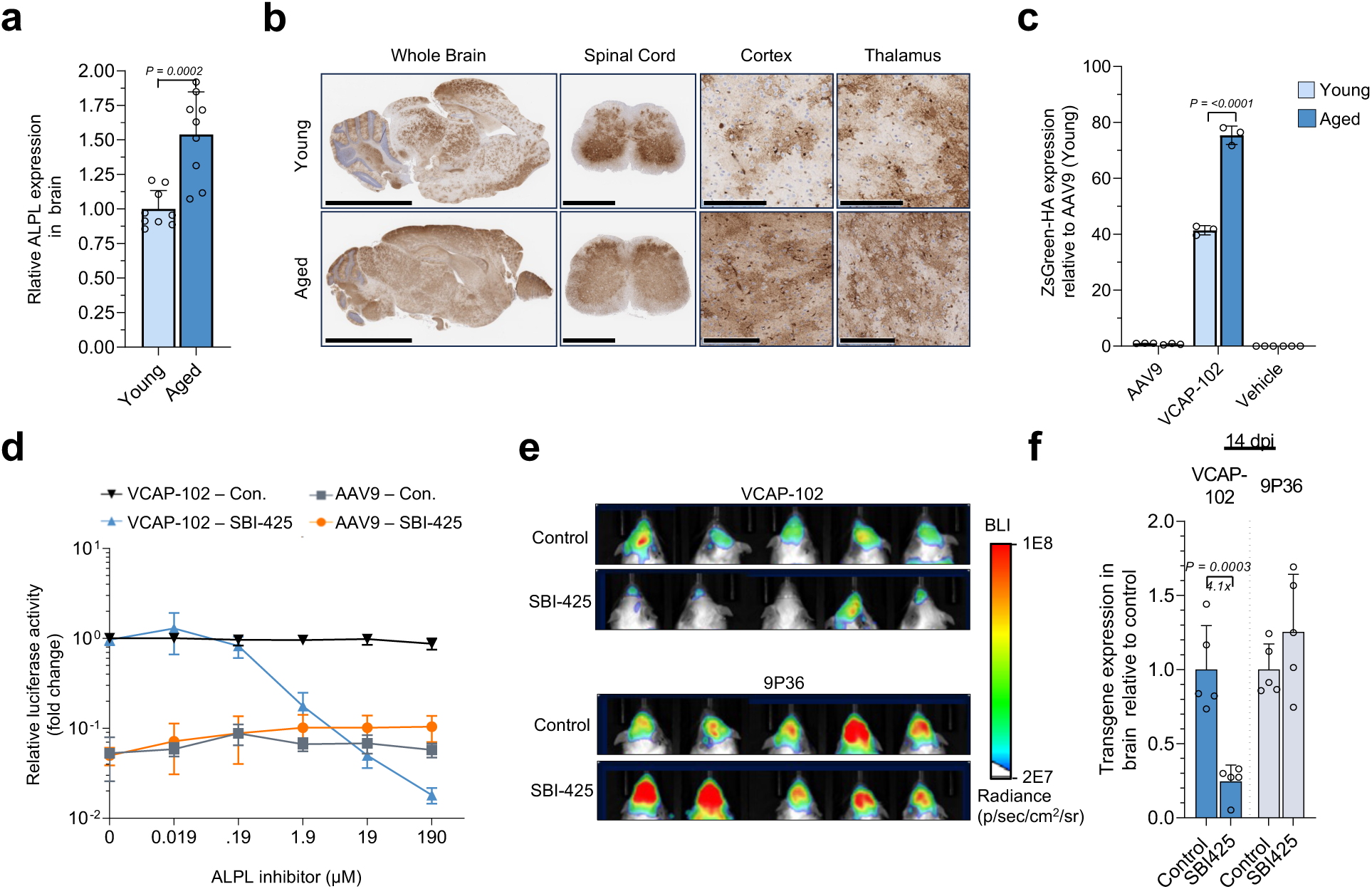
ALPL impacts VCAP-102 transduction *in vivo*. a, ALPL expression in young (6 week) and aged (18 months) mice relative to mTBP housekeeping gene. Values indicate mean ± SD (n = 9), normalized to young mouse data. **b,c**, CNS transduction by VCAP-102 in young and aged mice. **b**, Anti-HA staining 28d after transduction with 1E13 VG/kg VCAP-102 containing scAAV-CAG-zsGreen-HA. Scale bars: 5mm (whole brain), 1 mm (spinal cord), 200 µm (cortex, thalamus). **c**, Transgene mRNA expression in the brain of young and aged mice. Data show mean ± SD (n=3). **d**, HeLa cells were treated with ALPL inhibitor SBI-425 and exposed to 1E4 VG/cell AAV9 or VCAP-102-luciferase. Data (mean ± SD) is normalized to untreated cells. **e**, Brain transduction in mice treated with 35 mg/kg ALPL inhibitor SBI-425. VCAP-102 or 9P36 capsids with AkaLuc transgene were administered IV at 2E13 VG/kg. *In vivo* luminescence show luciferase expression in the brain 7 days after AAV administration. **f**, Brain transgene mRNA in mice from (**e**). Values indicate mean ± SD (n = 5) normalized to vehicle control. All *P* values are derived from an unpaired two-tailed t-test.

### *In silico* modeling of ALPL-Capsid interaction

In order to elucidate the molecular mechanism of capsid-ALPL interaction, *in silico* modeling of the VCAP-102-ALPL complex was performed with AlphaFold. AlphaFold is the current state-of-the-art protein structure prediction model, and it has been used in various peptide-protein binding prediction applications^36–38^. In agreement with our experimental data showing competitive binding with SBI-425, the AlphaFold-models predicted that the engineered VCAP-102 peptide occupies the catalytic pockets of the ALPL protein dimer, forming a depression adjacent to the central crown domain (Fig. 6a, Extended Data Fig. 8a). Consistent with our cross-species results, the ALPL residues predicted to form contacts with the capsid are conserved across human, cynomolgus macaque and mouse (not shown). AlphaFold modeling identified multiple residues on ALPL predicted to form contacts with the conserved SPH[+] motif of the VCAP-102 capsid (Fig. 6b). Strikingly, point mutation in those residues resulted in significant loss of transduction (data not shown). These observations not only validate the accuracy of the *in silico* model prediction, but it provides a potential molecular mechanism for the interaction of VCAP-102 capsid with a human endothelial receptor to gain access to the CNS.

**Fig. 6.**
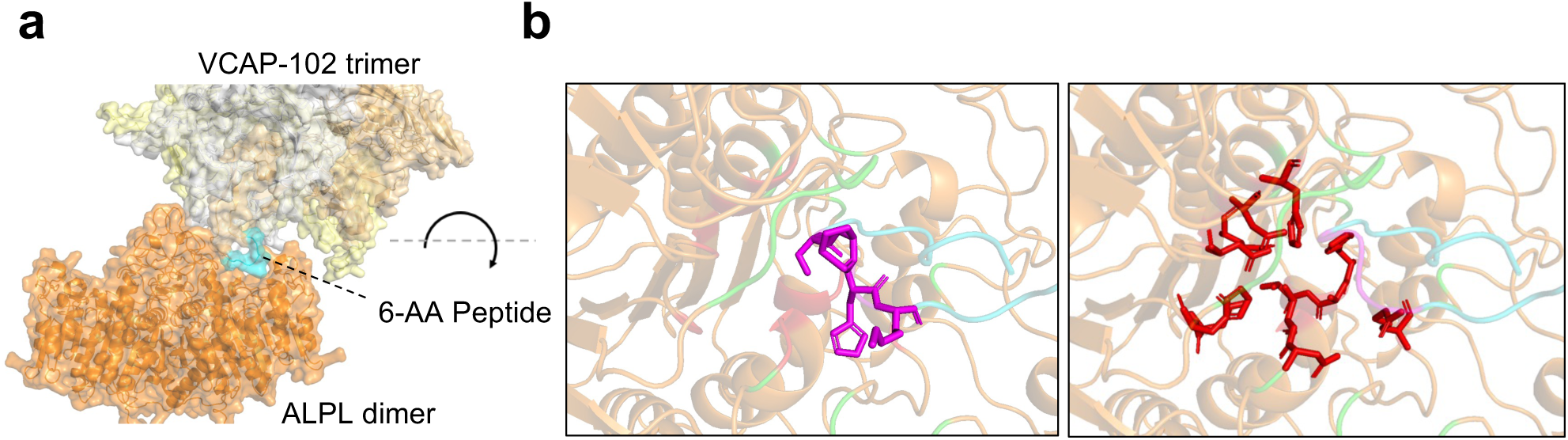
In silico prediction and validation of ALPL-VCAP-102 interaction. a, AlphaFold2-multimer predicted structure of ALPL dimer (orange) and VCAP-102 VP3 trimer (white, yellow, wheat) with VCAP-102 6-AA peptide shown in turquoise. **b,** Top-down view of the ALPL active pocket bound to the VCAP-102 peptide (turquoise). Left panel: conserved SPH[+] residues on the peptide are shown in magenta, ALPL residues predicted to interact with VCAP-102 are shown in green. Right panel: ALPL residues in contact with the SPH motif of the peptide are highlighted in red.

## Discussion

Here, we describe the use of RNA-driven AAV directed evolution to identify a bioengineered capsid variant, VCAP-102, with enhanced BBB penetration in primates and rodents. VCAP-102 was identified among a family of capsids sharing the minimal motif G453-XXSPHX[+]X-Q462. The fact that the same motif was recovered repeatedly from independent libraries suggests a near-exhaustive coverage of the sequence space which could result from our shorter 6-AA random insertion design. It is noticeable that no equivalent capsid was previously isolated from multiple AAV9 evolution screens using different surface loops^10,12,13,15^. This could suggest that structural diversification of capsid libraries can broaden the repertoire of receptors used by engineered AAV.

The VCAP-102 capsid displayed broad CNS transduction from intravenous dosing, and achieved gene transfer in 20-30% of SMI311-positive neurons and 60-77% Sox9-positive astrocytes in both mouse and primates at a dose of 1E13 VG/kg, which is less than 10% of the dose used with the AAV9-based Zolgensma^6,39^ in infants. The broad and robust CNS transduction at low dose, consistency of capsid tropism across species, and low liver transduction relative to AAV9 indicate that VCAP-102 could provide a markedly improved therapeutic window relative to AAV9 by achieving clinically effective gene transfer in the CNS with minimal liver toxicity.

We identified ALPL as the vascular receptor used to transport VCAP-102 across the BBB. Direct interaction of VCAP-102 with human ALPL was confirmed in a cell-free model and ALPL ectopic expression increased capsid transcytosis by >100-fold in a transwell model. ALPL is highly expressed in the brain vasculature^26,40,41^ and belongs to the glycosylphosphatidylinositol (GPI)-anchored (GPI-AP) family of membrane-anchored proteins. Strikingly, all three other receptors used by BBB-penetrant AAV capsids, namely Ly6A^18,19^, Ly6C1^20,21^ and CA4^21^, belong to the GPI-AP family. Several reasons could explain such preference. GPI-enriched endosomal compartments (GEEC) are the preferential trafficking route of natural AAV capsids^42^, and another major transcytosis route, caveolae-mediated vesicle trafficking, is suppressed in brain endothelium^43^. Together, this could explain why GPI-enriched vesicles remain as the most active transcytosis route to transport AAV and potentially other large cargos across the BBB.

ALPL is highly conserved across species, and we confirmed functional interaction of VCAP-102 with human, macaque, mouse and porcine ALPL *in vitro* (Fig. 4b). Not only does this suggest that VCAP-102 vectors will be effective in human patients, but this could also greatly facilitate testing of future brain-targeted therapeutics in common animal models with no need for surrogates or humanized transgenic animals. Another crucial benefit of identifying the mechanism of action is the potential to anticipate the level of gene delivery in the patient population based on receptor expression. We observed a strong correlation between VCAP-102 transduction and ALPL expression levels, as indicated by a strikingly high transduction in marmosets (Extended Data Fig. 5) and aged mice, which both express high levels of ALPL in the brain^26,44,45^.

VCAP-102 binds ALPL with an affinity of ∼20 nM at pH 7.4 and is rapidly released at pH 5. These properties are common amongst other known receptor ligands used for BBB transport, such as antibodies directed against transferrin receptor, CD98hc or IGF1R^29,46,47^. All these transport vehicles show affinities within the 10-30 nM range and rapid dissociation at low pH. This would suggest that efficient transcytosis across the BBB follows the same general rules regardless of the nature of the receptor or the trafficking pathway. Those observations raise the tantalizing possibility that ALPL could be used for BBB transport of non-viral delivery modalities such as antibodies, nanobodies, peptides or aptamers, similarly to other BBB receptors. Our work provides strong *in vivo* and mechanistic evidence that ALPL can be harnessed to transport large molecular complexes across the BBB with high efficiency and broad distribution, and that these properties can be faithfully recapitulated in rodents, primates and eventually humans.

## Methods

### AAV capsid libraries

AAV9 “Hotspot” Library was assembled through a two-step cloning process (Extended Data Fig. 1). First, primers containing 6 degenerate NNK codons inserted at each of 153 capsid surface positions, followed by a unique restriction site and a position-specific barcode, were used to generate a series of 5’ capsid fragment, that were cloned by Gibson assembly into matching CAG-driven TRACER™ vectors and transformed into supercompent E. coli. Resulting vector pools were linearized using the restriction site between the peptide insertion and the barcode, and the remaining 3’ capsid sequences were inserted by Gibson Assembly. To generate TRACER-NHP libraries, a series of 6-AA random peptides encoded by NNK codons was inserted into the VR-IV loop of a CAG-driven TRACER vector using Gibson Assembly. During later steps of *in vivo* selection, library cDNA recovered by RT-PCR from animal tissue (see library screening methods) were cloned into TRACER vectors containing a human synapsin 1 (hSyn1) promoter. The third-pass synthetic library containing 1,500 selected variants was synthesized as an oligo pool by Twist Biosciences (San Francisco, CA) using two divergent nucleotide sequences for each peptide variant^22^ and cloned into a TRACER-hSyn1 vector.

### Virus production

Virus production was performed in suspension HEK293T maintained in FreeStyle™ F17 Expression Medium supplemented with GlutaMAX, penicillin/streptomycin, and 0.1% Pluronic F-68 (all from Gibco). Libraries were generated by triple transfection with 1 mg pAdDeltaF6 adenovirus helper plasmid, 500 µg REP plasmid and 16 µg library DNA per 1 liter of culture. For recombinant viruses used in validation studies, REP-CAP plasmids containing various capsid candidates were used for co-transfection of suspension HEK293T together with pAdDeltaF6 adenovirus helper plasmid, transgene plasmid at a 2:4:1 ratio. Cells were harvested 72 h after transfection by low-speed centrifugation and lysed by addition of 0.1% Triton X-100 (Thermo Scientific) and treated with nuclease for 1 hour. Lysate was clarified, brought to 1M NaCl, and fractionated on two successive rounds of iodixanol gradients as previously described^48^. Buffer exchange was performed on Sartorius Vivaspin® 100K PES columns with phosphate buffered saline containing 200 mM total NaCl and 0.001% Pluronic F-68 (Gibco). The final virus samples were titered by digital droplet PCR (BioRad) using a REP-specific primer/probe set for libraries and a CMV primer/probe set for recombinant virus (Supplementary Table 2). Final virus preparations were tested by silver stain of PAGE gels and Endosafe endotoxin assay (Charles River Laboratories, Wilmington, MA, USA).

### In vitro AAV9 “Hotspot” Library Transduction Assay

AAV9 “Hotspot” DNA library was constructed and the viral library was produced via triple transfection in suspension HEK293T cells and a purified with a single iodixanol fractionation step. Virus was added to HEK293T at a MOI of 1e5 VG/cell. Cells were harvested 48 hours after transduction, RNA was isolated using an RNeasy kit (Qiagen) and reverse transcription was performed with the Quantitect kit (QIAGEN). Library cDNA was subjected to PCR for Illumina NGS sequencing as described below in the *in vivo* screening methods.

### Animals and Husbandry

C56Bl/6J and BALB/c mice (22–30 g; 7–8 weeks, or 72 weeks for aged mouse experiment) were purchased from The Jackson Laboratory (Bar Harbor, ME, USA). Animals were housed in a 12-h light:12-h dark environment and provided food and water ad libitum. All animal protocols were approved by the Voyager Therapeutics Institutional Animal Care and Use Committee (IACUC). For primate studies, animals were pre-screened for AAV9 neutralizing antibodies using a AAV9-luciferase-based assay. Upon dosing, all primates were housed individually in stainless steel cages, and provided a standard primate diet for the given species and provided continuous water. Animals were housed in a 12-hour light/dark cycle, an approximately 70°F atmosphere, and monitored cage-side daily. Animals were given additional supplements and various cage-enrichment devices. All procedures were in compliance with the Animal Welfare Act, the Guide for the Care and Use of Laboratory Animals and the Office of Laboratory Animal Welfare. All protocols were approved and monitored by the local IACUC.

### Library screening by *in vivo* selection

AAV libraries were injected into mice (n = 3) at a dose of 5e13 VG/kg into the lateral tail vein. Animals were euthanized 28 days postinjection and perfused with cold PBS. Brain, spinal cord, heart, and liver were snap-frozen in liquid nitrogen, and stored at -80°C. For primates, on the day of dosing animals were pre-treated with intramuscular injection of dexmedethomidine (15 ug/kg) and ketamine (10 mg/kg). Viral libraries were administered intravenously as a single saphenous vein infusion over the course of 30 minutes - 1 hour using a syringe pump. After a 28-day in-life period, euthanasia was performed using pentobarbital sodium intravenous injection (100 mg/kg) followed by bilateral thoracotomy. Animals underwent transcardial perfusion with ice-cold heparinized (100 U/mL) saline over the course of ten minutes. Brains were chilled in cold saline, sectioned into 3-4 mm coronal slices and snap frozen. All tissue samples were snap frozen in dry ice-cooled isopentane and stored at -80°C. Mouse and primate whole-brain RNA was extracted using trizol extraction, and mRNA was purified using Oligotex beads (QIAGEN). mRNA was treated with EZ-DNase (Life Technologies) and subjected to reverse transcription with a library-specific reverse primer using Superscript IV first-strand synthesis kit (Life Technologies). Capsid library cDNA was then amplified in multiple 50-µL PCR reactions unsing Q5 HotStart high-fidelity 2X master mix (NEB).

### NGS and bioinformatics analysis

NGS amplicon libraries were generated and analyzed as previously described^10,22^. Briefly, NGS primers containing indexed Illumina adapters fused to capsid-specific handles were used for low-cycle (<15 cycles) nested PCR amplification using recovered library amplicons as templates. The resulting fragments were gel-purified and subjected to Illumina Nextseq500 sequencing. NGS data was analyzed by a custom AAV amplicon-sequencing pipeline for enrichment analysis.

### *In vivo* characterization of TRACER AAV variants

Reporter plasmids containing zsGreen-HA, H2B-HA or H3F3-HA transgenes under control of the CAG promoter were generated using standard molecular cloning techniques. Capsid variants were injected into mice (n = 3) at a dose of 1e13 VG/kg (unless otherwise indicated in the text) via lateral tail vein. Animals were euthanized 28 days postinjection. One half of each organ was fixed in 10% neutral buffered formalin for immunostaining and the other half was snap frozen for DNA/RNA extraction. Capsid variants were injected into primates anesthetized with ketamine (10 mg/kg) at a dose of 1e13 VG/kg (African green monkeys) or 2e12 VG/kg per (marmosets) via saphenous vein infusion. Following sedation and euthanasia at day 28, peripheral tissues were collected using ∼1g of each tissue for snap freezing and ∼1 g for fixation in 10% NBF. Brains were hemisected, left brain hemisphere was sectioned in 3-4 mm coronal sections and flash frozen while the right hemisphere was sectioned and fixed by 10% NBF for histology and immunohistochemistry.

### Immunohistochemistry and imaging

Tissues were fixed in neutral buffered formalin (NBF) for 36-48 hours at room temperature, embedded into paraffin blocks, and sectioned at 5 µm thickness. The following human tissues were ordered from Biochain Institute: Cerebral Cortex (Cat # T2234042), Cerebellum (Cat # T2234039), Thalamus (Cat # T2234079) and Human Adult Normal Tissue, Array I of 64 Specimens (Cat #: T8234708-5).

#### Immunohistochemistry

Mouse tissue staining was conducted on the Leica Bond RXm platform using standard chromogenic methods. Slides were stained with anti-HA antibody (Cell Signaling #3724S, 1:800 dilution) followed by HRP-conjugated anti-rabbit secondary polymer and chromogenic visualization with diaminobenzidine (DAB). Hematoxylin counterstain was used to label nuclei. African green monkey HA tag staining was performed on a Ventana Discovery. FFPE sections were incubated with anti-HA antibody (1:400 dilution) followed by incubation with anti-rabbit-HQ (Roche; Cat#. 760-4815) and anti-HQ HRP (Roche; Cat#760-4820). IHC signal was detected using the Discovery Chromomap DAB kit (Cat#. 760-159). Slides were scanned on a Leica Aperio Versa 200 and imaged at 5X, 20X, and 40X magnification.

Marmoset stainings (HA and Myc tag) and all ALPL stainings were performed on a BondRx with BOND Polymer Refine Detection kit (Catalogue No. DS9800). Antigen retrieval was carried out with TRIS buffer at pH 9 for 20 mins at 95°C using Bond Epitope Retrieval Solution 2 (Leica Biosystems). HA and Myc tag wer detected using anti-HA (see above) an anti-Myc monoclonal antibody (Cell Signaling Technology; Cat# 92013S) at 0.035µg/mL. ALPL was detected with Anti-ALPL monoclonal antibody (Invitrogen; Cat# MA5-17030) at 1:10,000 dilution. Slides were scanned on a Leica Aperio Versa.

#### Immunofluorescence

Co-staining of Sox9 or NeuN with HA in FFPE mouse tissues was conducted on a Leica Bond RX platform. The following antibodies were used: anti-Sox9 (ab185966, Abcam, 1:1,000:), anti-NeuN (ab177487, Abcam, 1:1000), anti-HA (3724, Cell Signaling, 1:800). A two-step sequential staining method was used. The first antibody was added and detected using an HRP-conjugated anti-rabbit secondary polymer followed with opal 520 (for Sox9) or opal 620 (for HA) reagent tyramide signal amplification. The first antibody was stripped and the next primary antibody was added for 30 minutes and detected using an HRP conjugated anti-rabbit secondary polymer, followed with reciprocal opal 620 or opal 520 reagent tyramide signal amplification. Nuclei were visualized with spectral DAPI (Akoya Biosciences). Co-staining of SMI311 and GFAP in African green monkey tissues was conducted on a Ventana Discovery Ultra.

Primary antibody incubation with rabbit anti-HA antibody was followed by incubation with OmniMap anti-rabbit HRP (05269679001, Roche Diagnostics). Fluorescence signal was detected with FITC fluorophore (07259212001, Roche Diagnostics). Sequentially, neurons were labelled with SMI-311 (837801, BioLegend) and detected using Goat anti-mouse Alexa Flour 647 and astrocytes were labelled with GFAP Alexa Flour 555 (3656S, Cell Signaling Technology). Slides were mounted with Prolong Diamond (P36961, Invitrogen) and were imaged on the Olympus VS200 microscope.

#### Co-localization analysis

Images were loaded into HALO Image Analysis Software (Indica Labs). Quantification of chromogenic HA-positive cells or colocalization of HA tag with cell-specific markers in mouse and primate brains were performed with Indica Labs HighPlex or Multiplex algorithms. Nuclear segmentation classifiers were trained to distinguish between positive and negative cells.

### Vector mRNA and DNA quantification in mouse tissue

For transgene mRNA quantification, total RNA was extracted from 100–200 mg of tissue using RNeasy plus universal kit (QIAGEN), and reverse transcription was performed with Quantitect kit (QIAGEN). Spliced transcripts were quantified by TaqMan^TM^ PCR using GloX4 primer-probe set specific for transgene exon-exon junction (Table 2). Murine TATA box-binding protein (mTBP) RNA was used as a housekeeping reference. Relative expression levels were calculated from the DCt values and normalized to AAV9 samples. For vector DNA quantification, 20 mg tissue was processed using the Blood and Tissue DNeasy kit (QIAGEN). 100 ng DNA was used for TaqMan^TM^ PCR quantification with transgene-specific probe/primer sets normalized to the TERT gene (Supplementary Table 2).

### Vector mRNA and DNA quantification in primate tissue

Tissue samples (2-mm biopsy punch) were homogenized in Quantigene Lysis Mixture (Thermo Fisher Cat. #QP0523) with 2% (v/v) Proteinase K (Thermo Fisher Cat. #QG0517), mechanically disrupted using a Geno-Grinder® (SPEX Sample Prep), and incubated 1h at 65°C. Lysates were clarified by centrifugation and split into two fractions for RNA and DNA extraction. RNA was extracted with Qiagen RNeasy Universal Mini Kit (Cat. #73404) following manufacturer’s instructions. DNA was extracted using the Qiagen DNeasy Blood and Tissue Kit (Cat. #65906). Total RNA was digested with DNase I (Sigma Aldrich Cat. #AMPD1-1KT) and reverse transcribed using High-Capacity cDNA Reverse Transcription Kit (Thermo Fisher, Applied Biosystems, Cat. #4368813). Transgene expression was measured either by RT-qPCR (Bio-Rad CFX384) or by RT-ddPCR (digital droplet PCR, Bio-Rad QX200 system) in duplex reactions using a transgene-specific primer probe set (Supplementary Table 2) normalized to rhTBP housekeeping gene.

Vector DNA was measured by duplex ddPCR reactions on whole cell DNA with Taqman primer-probe sets specific to the CMV promoter (African Green monkey experiments) or to the HA or Myc epitope tag (marmoset experiments). RNaseP was used as a genomic copy number control (Supplementary Table 2).

### Retrogenix Cell Microarray

The Retrogenix cell microarray was performed by Charles River Laboratories. cDNA expression vectors encoding over 6000 human membrane proteins, secreted or cell surface tethered-secreted proteins were arrayed across a series of slides in duplicate and overlaid with HEK293 as previously described^49^. Cells were added to the slides for reverse transfection and expression of the specific cDNA spotted at that location.

Following cell fixation, AAV9 and VCAP-101 were added to each slide at an MOI of 3E5 particles/cell. Bound AAV was detected with a 1:500 dilution of anti-AAV9 antibody (clone HL2372, Millipore Sigma, MAB2309) followed by AF647 anti-mIgG H+L secondary antibody. Slides were imaged using ImageQuant and candidate receptors were identified as duplicate spots showing a raised signal compared to background levels. A combined 27 interactions were identified in the library screen. A confirmation screen was carried against the 27 candidates as described above with AAV9, VCAP-101 and additional controls.

### Human ORFeome Screen

A stable HEK293T cell pool was established using a lentiviral human ORFeome library (Millipore Sigma, ORFPOOLWG) containing 17,000 human cDNAs, covering 14,000 genes. Following lentiviral transduction, selection was carried out by maintaining cells in complete DMEM supplemented with 1.5 µg/mL puromycin. For the first two rounds of screening the HEK293T ORFeome cell pool was transduced with 1E4 VG/cell of VCAP-102-GFP. GFP(+) cells were sorted on a BD FACSmelody and expanded before the next round of transduction and sorting. In the third and final round of screening, cells were co-transduced with 1E4 VG/cell of VCAP-102-GFP and 1E4 VG/cell AAV9-mCherry in order to differentiate between cellular factors involved in general AAV9 transduction (GFP(+)/mCherry(+)) and those specific to VCAP-102 transduction (GFP(+)/mCherry(-)). The 24-nucleotide barcode situated in the 3’UTR of each ORF was used for NGS enrichment analysis after each cycle. DNA was extracted from 1-5E5 sorted cells using the DNeasy blood and tissue kit (Qiagen, 69504) and used for barcode amplification and NGS analysis.

### Cells and Expression Plasmids

HEK-293T and HeLa cells were maintained in Dulbecco’s modified Eagle medium (DMEM) (Gibco, 1056910) supplemented with 10% heat inactivated Fetal Bovine Serum (FBS) (Corning, 35-015-CV) and penicillin-streptomycin (Gibco, 15140122). Madin-Darby Canine Kidney (MDCK) cells were obtained from Millipore-Sigma (85011435) and maintained in EMEM media (ATCC, 30-2003) containing 10% heat-inactivated FBS (Corning, 35-015-CV). HEK293T-ALPL, HeLa-ALPL and MDCK-ALPL stable cell lines were generated with lentiviral vector from OriGene (RC205692L3V) and maintained in the media described above supplemented with either 1.5ug/mL (HEK-293T and HeLa) or 10µg/mL puromycin (MDCK) (Mirus Bio, MIR 5940). A clonal MDCK-ALPL line with high transcytosis activity (MDCK-ALPL clone 21) was isolated by limited dilution. Expression plasmids for AAVR, ALPL (SC320807), ALPL-V2 (SC323118), and mouse ALPL (MC228161) were purchased from OriGene. Cynomolgus Macaque (XM_005544525) and Porcine (XM_021097682) ALPL expression plasmids were cloned into the EcoRI and XbaI restriction sites of the PCMV6-AC (Origene) using gBlocks (Integrated DNA Technologies) encoding the gene.

### Transfection, siRNA, Transduction Assays

HEK293T cells grown in 96-well plates were transfected by calcium phosphate precipitation. AAV was added 24h post-transfection at an MOI of 1E4 VG/cell. Several transgene reporter constructs were utilized in transduction assays: NLS-EGFP-WPRE, NLS-mCherry-WPRE and Firefly Luciferase Luc2-T2A-GFP, all cloned in single-stranded ITR AAV plasmids under a ubiquitous CAG promoter. 24 hours post-transduction, cells were visualized on an Eclipse Ti-S (Nikon) fluorescence microscope to visualize GFP or mCherry expression. Firefly luciferase activity was quantified using the ONE-Glo™ EX Luciferase Assay System (Promega, E8130) and read with the Synergy NEO2 multimode reader (BioTek). HeLa cells were transfected with siRNAs using Lipofectamine 2000 (Invitrogen). 48 hours post transfection the cells were transduced with AAV at an MOI of 1E4 VG/cell. Fluorescence and luciferase activity were measured 24h post-transduction as described above. All siRNAs were purchased from Ambion.

### ALPL inhibition in cultured cells

For *in vitro* ALPL inhibitor assays the small molecule ALPL inhibitor SBI-425 (Millipore Sigma, SML2935) was resuspended in 1mL of dimethyl sulfoxide (DMSO) (Sigma-Aldrich) for a final concentration of 25mg/ml. The inhibitor and vehicle (DMSO) were diluted in DMEM supplemented with 10% FBS. After 1 hour at 37°C cells were transduced with AAV9 or VCAP-102 containing a CAG-Luc2-T2A-GFP transgene at an MOI of 1E4 VG/cell. After 4 hours, the media was removed and replaced with fresh complete DMEM. Luciferase activity was quantified 24h post-transduction.

### ALPL inhibition in mice

For *in vivo* administration of SBI-425, a 25 mg/mL stock solution was diluted in vehicle solution and injected peritoneally to female B6 albino mice (B6(Cg)-Tyrc-2J/J) at 35 mg/kg daily for four days. On the second day of SBI-425 dosing, mice were dosed intravenously with 2E13 vg/kg of VAP-102 or 9P36 harboring an CAG-EGFP-T2A-AkaLuc transgene. On day 7 post-AAV administration, transduction was measured in live animals utlizing the Akaluciferase-akalumine reporter system^50^ optimized for deep tissue penetration of the near-infrared emitted light. Prior to imaging, mice were injected intraperitoneally with 28 mg/kg of the Akaluciferase^50^ substrate Akalumine-HCl (TokeOni) (Sigma, 808350) formulated at 16.5 mM in water. Animals were then immediately anesthetized with isofluorane gas and imaged using an AmiHT imager (Spectral Instruments) to capture bioluminescence emission. Photon emission data were normalized as photon per second per square centimeter per steradian. Images shown were exported from the Aura imaging software, keeping the color scale identical for each individual image. Mice were sacrificed on day 14 post-AAV administration as described previously and tissues were processed for transgene RNA quantification.

### Antibody competition assay

HeLa cells were plated in a 96 well format. Human ALPL Antibody (Biotechne, MAB1448) and Mouse IgG1 Isotype Control (Biotechne, MAB002) were resuspended to 500 ug/mL in PBS, diluted in serum-free DMEM and added to cells for 1 hour on ice. Following the incubation, cells were transduced with AAV9 or VCAP-102 containing a CAG-Luc2-T2A-GFP transgene at an MOI of 1E4 VG/cell for 4 hours at 37°C. After 4 hours media was removed and replaced with complete DMEM. Firefly luciferase activity was measured 24 hours post-transduction.

### PI-PLC and neuraminidase treatment

Phospholipase C Protein, Phosphatidylinositol-Specific (PI-PLC) from Bacillus cereus (Invitrogen, P6466) was diluted in serum-free DMEM. HeLa cells plated in 96-well format were washed in serum-free DMEM and treated with PI-PLC at varying concentrations. Cells were incubated at 37°C for 1.5 hours before washing and addition of 1E4 VG/cell AAV9 or VCAP102 harboring a Luc2-T2A-GFP transgene. The virus and cells were incubated at 37°C for 3 hours. After 3 hours, the media was removed and replaced with complete DMEM. Luciferase assay was performed 24 hours post-transduction. For neuraminidase treatment, HeLa cells were treated with *Vibrio cholerae* Neuraminidase (Sigma, N7885) diluted in serum-free DMEM for 1hr at 37°C prior to transduction. Cells were then washed and transduced with 1E4 VG/cell AAV9 or VCAP-102 containing a luciferase transgene. Luciferase assay was performed 24hr after transduction.

### Surface Plasmon Resonance

Binding of VCAP-102 and AAV9 to human ALPL (hALPL) was measured by Surface Plasmon Resonance (SPR) on Biacore 8K instrument. The capsids were immobilized on CM5 sensor chip (Cytiva, 29149603) by amine coupling and His-tagged hALPL (Sino Biological, 10440-H08H) was injected at 15.6 -250 nM concentration to measure binding kinetics. The surface was regenerated by injecting 10 mM glycine pH 1.5 for 30 seconds. The sensorgram was fitted to 1:1 binding model to calculate binding affinity. Alternatively, binding was confirmed in a reversed assay format, where His-tagged hALPL was captured via anti-His antibody pre-immobilized on sensor chip before injecting 0.0625-1 nM of capsid. The surface was regenerated by injecting two cycles of 10 mM glycine pH 1.7 for 30 seconds. A flow rate of 30 ul/min was used for SPR experiments and the sample and running buffer used was PBS-P+. pH-dependent binding of VCAP-102 to hALPL was measured by SPR on Biacore 8k instrument. The capsid was immobilized on CM5 sensor chip by amine coupling and His-tagged hALPL was injected at 7.8 -250 nM concentration at pH 7.4 using A-B-A injection mode. The dissociation was performed at pH 7.4 (PBS-P+ buffer) or pH 5.5 (20 mM MES, 150 mM NaCl, 0.05% Tween 20 buffer). The surface was regenerated by injecting 10 mM glycine pH 1.5 for 30 seconds.

### Transwell assay

MDCK or MDCK-ALPL clone 21 cells (see description above) were plated at a density of 200,000 cells in 250 μl of complete growth media to the apical side of a 24-well Transwell^®^ insert (Corning, 3495).

Complete growth medium (1 ml) was added to the basolateral chamber and plates were incubated for 2-3 days to allow polarization and barrier formation. Electrical resistance was measured using an EVOM Epithelial Voltohmmeter to verify the integrity of tight junctions. The media in the apical chamber was replaced with 250 μl of media containing AAV9 or VCAP-102 with matching transgenes, and the media in the basolateral chamber was replaced with 600 ul of fresh DMEM. For ALPL inhibitor assay, TNAPi (EMD Millipore, 613810) was added to the top chamber during AAV transduction. Following a 24-hour incubation, MDCK cells and media from both chambers were collected separately. AAV particles were quantified by qPCR with a Taqman primer/probe set specific for the CMV enhancer. The viruses in the bottom chamber were also detected by dot blot on a nitrocellulose membrane with a biotinylated anti-AAV9 antibody (Thermo Fisher, 7103332100).

### Cryo-EM data collection and image processing

AAV sample was applied to glow discharged graphene oxide grid. Grids were plunge frozen using a Vitrobot Mark IV (FEI, Inc.). Data collection was carried out using a Thermo Fisher Scientific (Hillsboro, Oregon) Glacios Cryo Transmission Electron Microscope operated at 200kV and equipped with a Falcon 4 direct electron detector. Automated data-collection was performed with Leginon software at a nominal magnification of 150,000x, corresponding to a pixel size of 0.923 Å. A total of 2,359 movies were recorded with a total exposure of 19.59 e^−^/Å^2^. All the subsequent image processing was performed in cryoSPARC 3.3. A total of 61k particles were picked for sample AAV using blob picker. These particles were extracted from the cryoTEM images and subjected to three rounds of reference-free 2D classification. The best 10k particles were used for the subsequent 3D reconstruction. The final 3D reconstruction was generated after ab-initio and homogenous refinement with icosahedral symmetry imposed. The resolution of the final 3D reconstructions was determined by the Fourier shell correlation (FSC) between two independent half maps, which was 3.4 Å at FSC = 0.143.

### Structure modeling

Prediction of the receptor-peptide complex was performed using AlphaFold2-multimer using each of the 5 model parameters provided^51,52^. MSA generation and structural prediction were performed using slightly modified ColabFold scripts to enable local execution^52^. We used multiple seeds to generate ensembles of the peptide-receptor complex for each model parameter. The side chains are then remodeled using AttnPacker^53^. Each of the complex was then scored using Arpeggio^54^ to extract the interacting residues between our predicted peptide structure and X-Ray diffraction structure [PDB: 7YIV]^55^. We counted the number of times a pair of residues was interacting with each other between the peptide and the receptor across the ensembles to generate a heatmap of probabilities for interacting pairs of receptors. 3D structure images were generated using PyMol.

### Constrained docking

We docked the full VP3 structure of our capsid onto the ALPL receptor structure with the constraints that the interaction pairs of residues identified using our AlphaFold-multimer predicted structure. The docking procedure was performed using local implementation ColabDock^56^.

### Statistical analysis

Unpaired two-tailed t tests were performed in Graphpad Prism and are reported in figures or figure legends. A p value < 0.05 was considered significant. Results are reported as mean ± SD.

## Contributions

M.N., T.C.M. and B.H. conceived and designed the experiments.

T.C.M., B.H., I. S., X-Q. R., T. K., E. R., K. G. performed library screening and receptor-related experiments.

J. Lu., J.Li. and W. W. performed next-generation sequencing analysis.

W. C. and J. L. performed in silico modeling of receptor-capsid interactions.

N. P., H. K., A. S. K., C. E. performed immunohistochemistry, immunofluorescence and image analysis.

J. T., J. C., A. B. and D. M., performed quantitative tissue transduction analysis. M.A.C, R. K. and Z. T., performed AAV library and recombinant AAV production.

C.H-Y.C., A. S-Z., A. S. K., A. J. and J. Y. organized and executed the mouse and primate *in vivo* studies.

## Acknowledgements

This study was funded by Voyager Therapeutics. The authors thank Kyle Grant, Tim Fiore and Dillon Kavanagh for their help with virus analytics. Cryo-EM data collection and image processing were performed at Nanoimaging Services, Inc. Figures were created using images from biorender.com.

## Supplementary data

**Extended Data Fig. 1.**
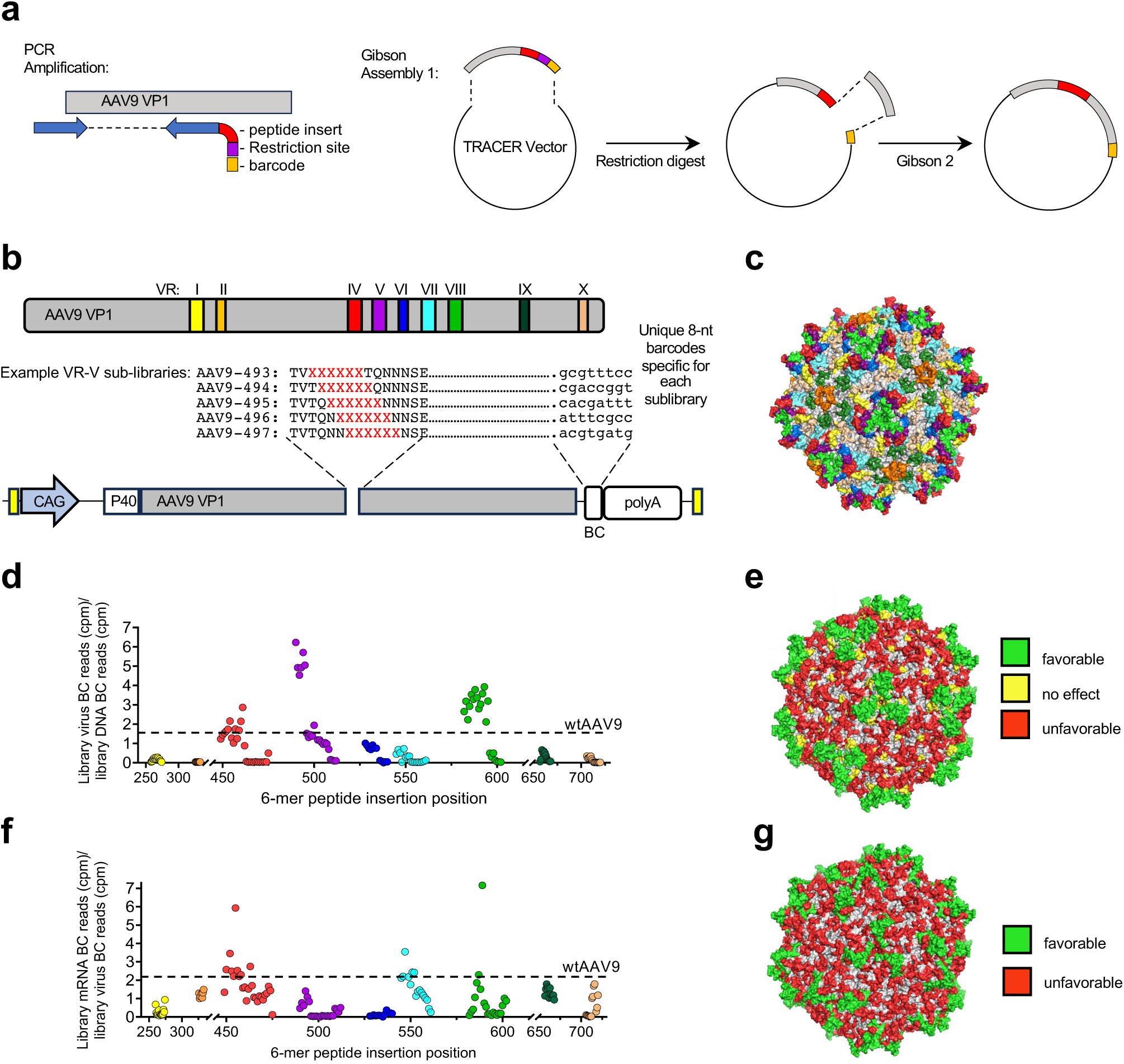
AAV9 “Hotspot” library reveals Variable Region IV as amenable to peptide insertion. **a**, Design of a barcoded AAV9 “hotspot” library. 153 Individual primers with a random 6-mer peptide inserted at each capsid surface position were used in a two-step cloning reaction to associate each insertion site to a unique 3’-end barcode (see Materials and Methods). **b**, Full length CAP gene is shown with the 9 variable regions (VRs) diagrammed. An example showing the VR-V insertion scan is shown. All DNA libraries were pooled together to generate a virus library by triple transfection. **c**, Structure of the AAV9 capsid (PDB: 3UX1) showing all positions included in the library, colored according to the VR loops shown in (**b**). **d**, Capsid viability score of each peptide insertion site, represented by the ratio of NGS barcode reads in the virus progeny relative to barcode reads in the DNA library. Each point represents the average data of n=3 virus production runs. AAV9 score is represented by a dotted line. **e**, Capsid viability scores overlayed on AAV9 structure indicating peptide insertion positions with positive (green), neutral (yellow) or negative (red) impact on virus production. **f**, Capsid transduction score representing the ratio of NGS barcode reads in RNA recovered from transduced HEK-293 cells relative to barcode reads from the viral library. Each point represents the average data of three separate virus productions. AAV9 is represented by a dotted line. **g**, Structural overlay of capsid transduction scores indicating peptide insertion positions with positive (green) or negative (red) impact on virus transduction.

**Extended Data Fig. 2.**
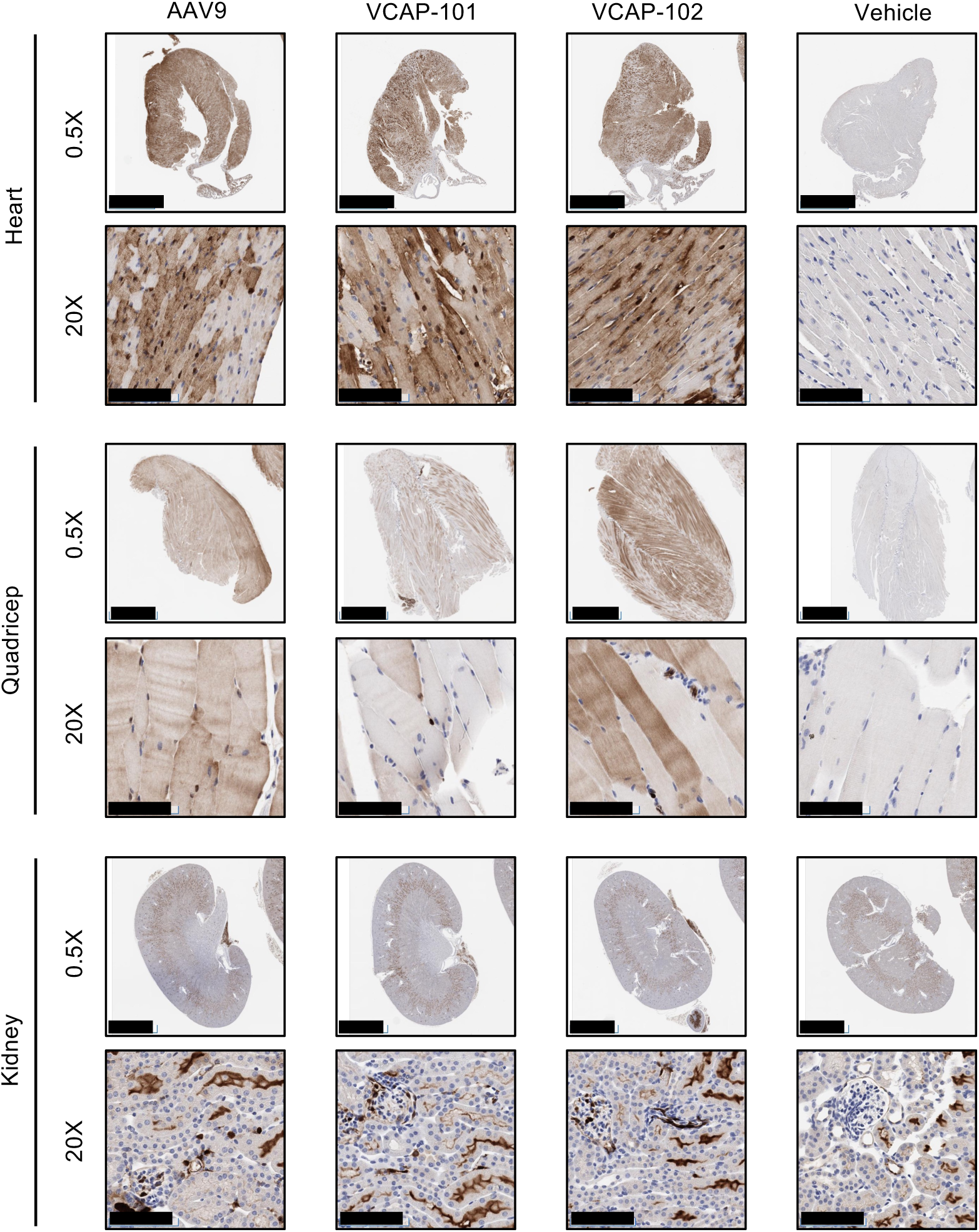
VCAP-101 and VCAP-102 peripheral tissue tropism in mice. HA tag was detected by IHC from FFPE sections of 12-week old BALB/c mice 28 days after IV injection of indicated capsids containing a self-complementary CAG-ZsGreen-HA transgene. Scale bars for 0.5X images represent 2.5 mm. Scale bars for 20X images represent 100 µm.

**Extended Data Fig. 3.**
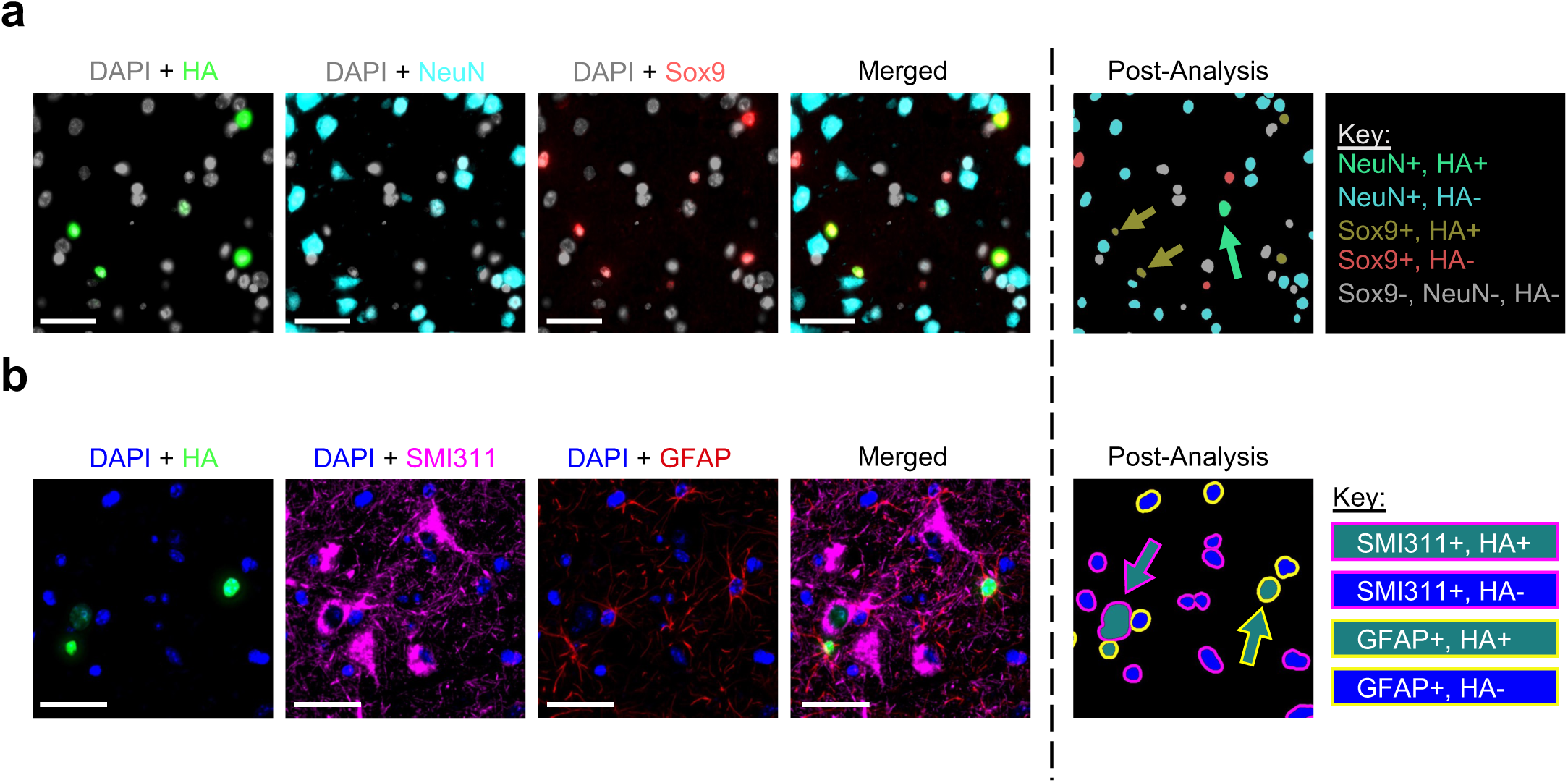
VCAP-102 cell type-specific tropism in African green monkeys. a,b, Indicated markers were detected by immunofluorescence from FFPE sections of the temporal cortex (**a**) or thalamus (**b**) of African green monkeys 28 days after IV injection of VCAP-102 containing a CAG promoter-driven, self-complementary H2B-HA transgene. Images were subjected to co-localization analysis using HALO imaging software (see methods). Scale bars represent 40 µm.

**Extended Data Fig. 4.**
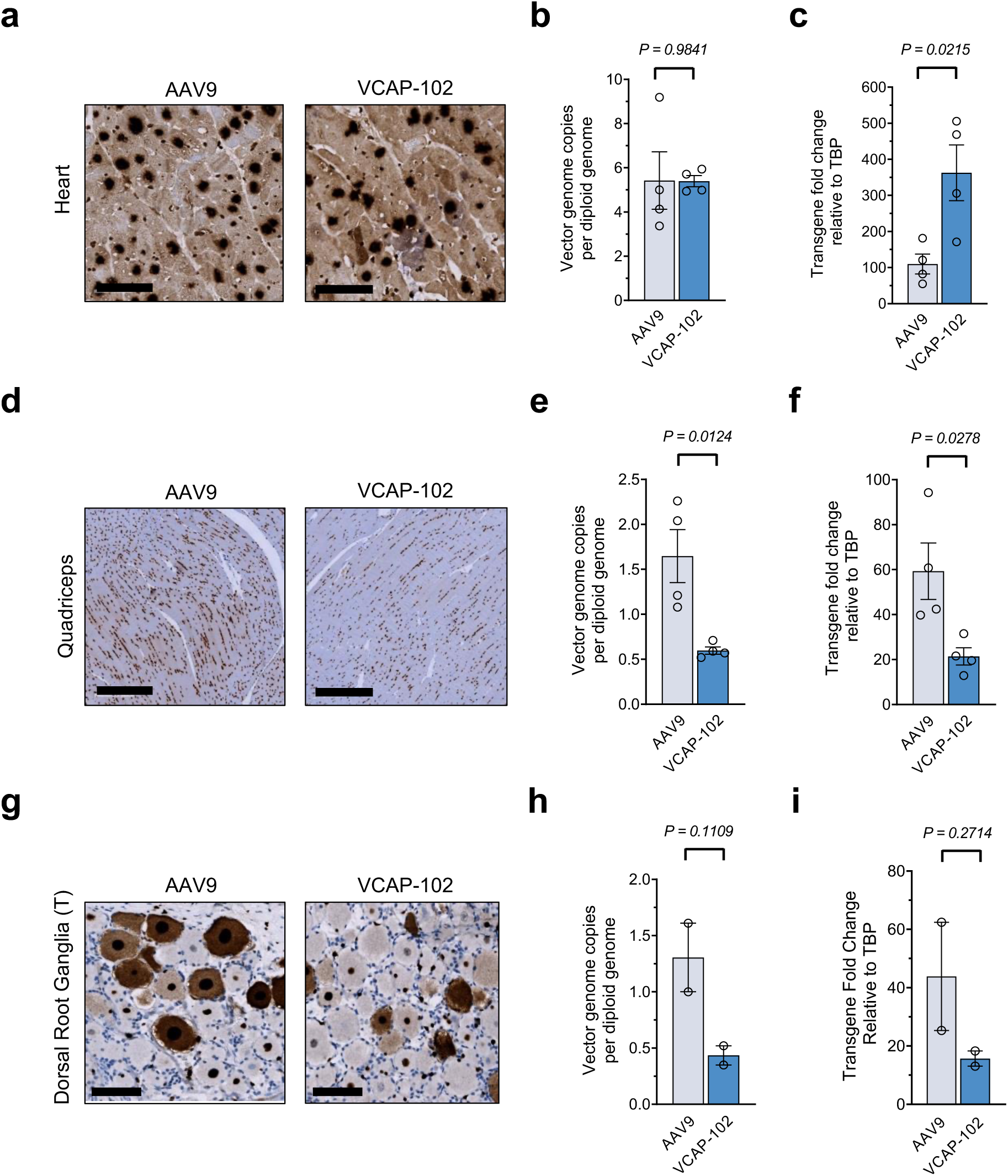
VCAP-102 tropism in non-CNS tissue in African green monkeys. Transgene expression was detected by IHC with an anti-HA antibody in sections from (**a**) heart ventricles, (**d**) quadriceps and (**g**) Dorsal Root Ganglia (DRG) of African green monkeys 28 days after i.v. injection of indicated capsid containing a self-complementary CAG-cH2B-HA transgene. Scale bars represent 60 µm. **b**, **e**, **h**, AAV genomes measured by ddPCR (n=4; 2 biopsy punches measured per monkey (n=2)). **c**, **f**, **i**, Transgene mRNA expression measured by qRT-PCR and normalized to TATA-binding Protein (TBP) (n=4; 2 biopsy punches measured per monkey (n=2)). **b**, **c**, **e**, **f**, **h**, **i**, Values indicate mean ± SD. *P* values were determined by unpaired two-tailed *t*-test.

**Extended Data Fig. 5.**
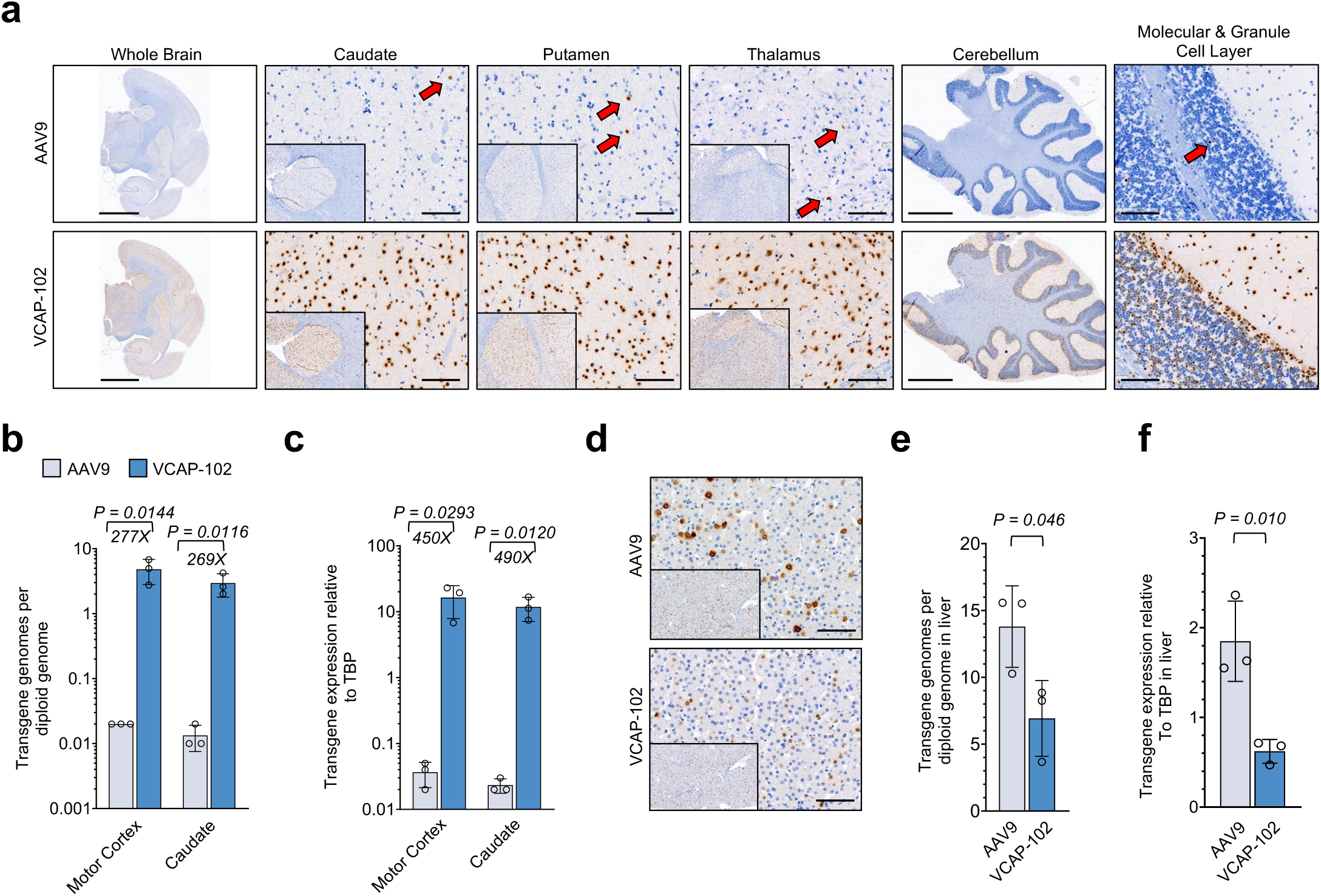
VCAP-102 displays increased CNS tropism in the marmoset brain. **a**, HA- or Myc-tagged transgenes detected by IHC from FFPE sections of marmosets (n=3) 28 days after IV co-injection of 2e12 VG/kg AAV9 (containing a self-complementary CAG-H2B-HA transgene) and 2e12 VG/kg VCAP-102 (containing a self-complementary CAG-H2B-Myc transgene). Scale bars represent 4 mm for brain, 2 mm for cerebellum, and 100 µm for others. Red arrows in the AAV9 images point to HA-positive nuclei. **b**, AAV genome quantification in indicated brain regions as measured by ddPCR (n=3). Values indicate mean ± SD. *P* values, unpaired two-tailed *t*-test. **c**, Real-time RT-PCR analysis of H2B-HA or H2B-Myc mRNA expression in the indicated brain region relative to marmoset TATA-binding Protein (marmTBP) (n=3). Values indicate mean ± SD. *P* values, unpaired two-tailed *t*-test. **d**, HA or Myc was detected by IHC from FFPE sections of marmoset livers as described in (**a**). Scale bars represent 100 µm. **e**, AAV genome quantification in liver as measured by ddPCR (n=3). Values indicate mean ± SD. *P* values, unpaired two-tailed *t*-test. **f**, Real-time RT-PCR analysis of H2B-HA or H2B-Myc mRNA expression in liver relative to marmoset TATA-binding Protein (marmTBP) (n=3). Values indicate mean +/- SD. *P* values, unpaired two-tailed *t*-test.

**Extended Data Fig. 6.**
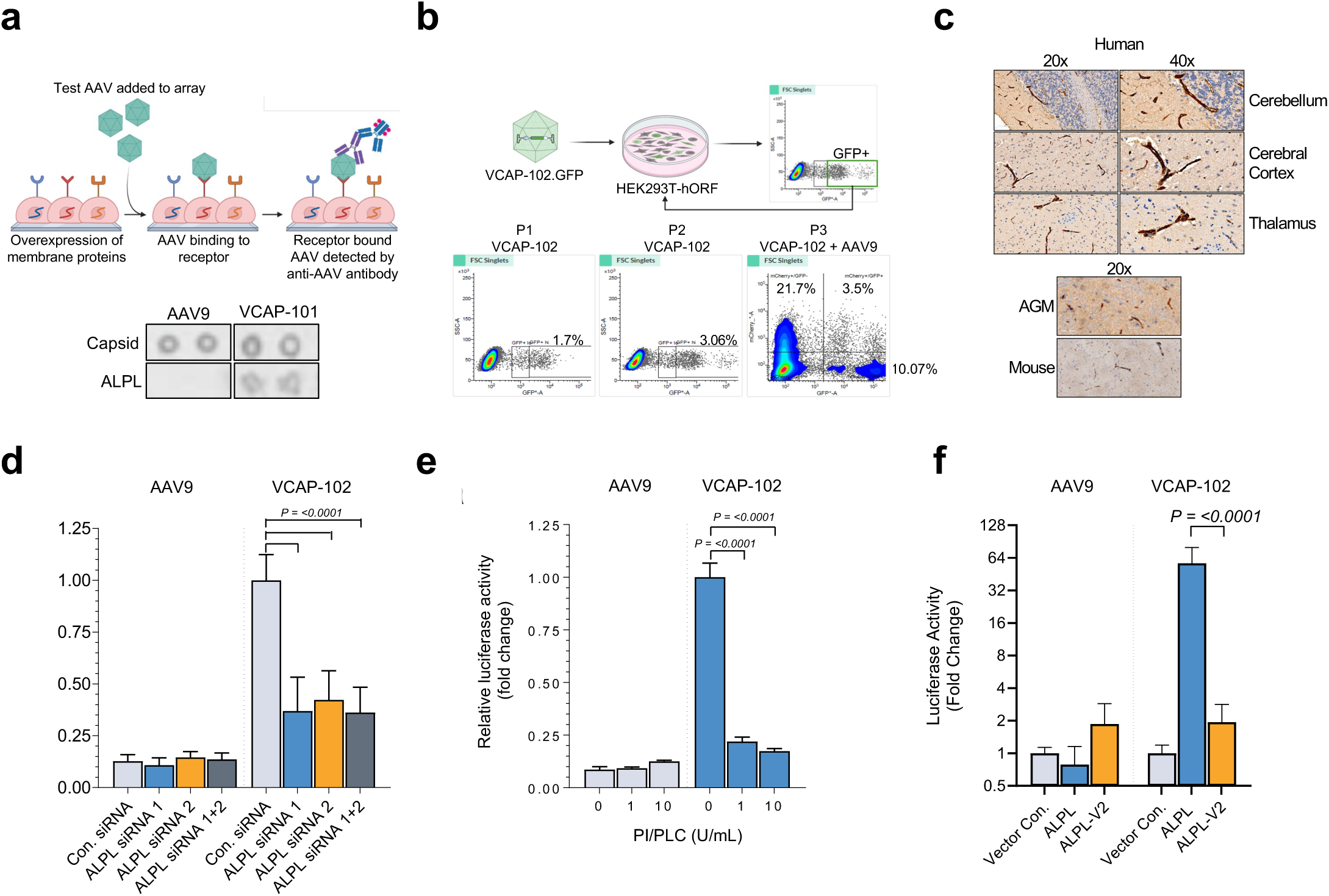
Identification of ALPL as a cell surface attachment receptor for VCAP-102. a, Retrogenix microarray screen. Expression vectors encoding >6000 human membrane proteins were arrayed on slides and overlaid with HEK293 cells for reverse transfection. AAV was added and cell-bound capsids were detected with an anti-AAV9 antibody. Bottom panel: Image from microarray screen demonstrating detection of virus spotted in gelatin (positive control) and VCAP-101 bound to cells expressing ALPL. **b**, Human ORFeome screen protocol. A lentiviral library containing 17,000 human ORFs was used to generate a stable HEK293T pool that was subjected to iterative rounds of transduction with VCAP-102-EGFP followed by sorting of GFP(+) cells. AAV9-mCherry was added to the third round of screening to identify EGFP(+)/mCherry(-) cells expressing a VCAP-102-specific receptor. Bottom panel: FACS gating strategy showing stepwise enrichment of permissive GFP(+) cells. **c,** Detection of ALPL by immunohistochemistry (IHC) in brain sections from human, African green monkey and mouse**. d**, Impact of ALPL depletion on VCAP-102 transduction. HeLa cells were transfected with siRNAs against ALPL and transduced with AAV9 or VCAP-102 containing a luciferase transgene. Luciferase data was normalized to VCAP-102 control siRNA (mean ± SD, n=3). **e**, Removal of cell surface GPI-AP proteins reduces VCAP-102 transduction. HeLa cells were treated with PI-PLC and transduced with AAV9 or VCAP-102 containing a luciferase transgene. Luciferase data (mean ± SD, n=3) is normalized to untreated controls. **f**, Plasma membrane localization of ALPL is necessary for VCAP-102 transduction. HEK293T cells were transfected with plasmid encoding an ALPL isoform defective for plasma membrane trafficking (ALPL-V2) and transduced with AAV9 or VCAP-102 encoding luciferase. 24h post-transduction a luciferase assay was performed. Data (mean ± SD, n=3) was normalized to vector control.

**Extended Data Fig. 7.**
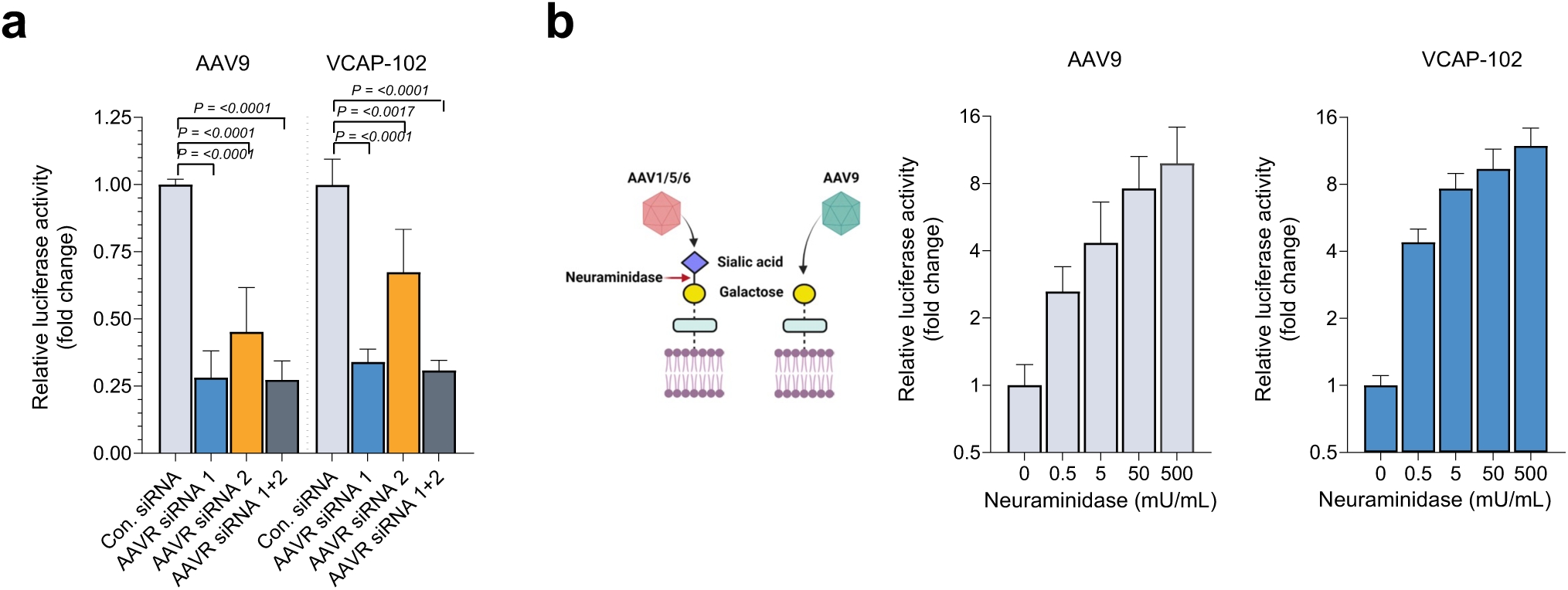
Impact of AAVR and galactose on VCAP-102 transduction. a, HeLa cells were transfected with three independent siRNAs against AAVR (KIAA0319L) or a scrambled control siRNA. 48h after transfection cells were treated with 1E4 vg/cell AAV9 or VCAP-102 expressing luciferase. Luciferase activity was analyzed 24h post-transduction and the data (mean ± SD, n=3) is normalized to control siRNA-treated cells . **b**, Galactose usage by VCAP-102 and AAV9. Left: mechanism of N-terminal sialic acid cleavage by neuraminidase resulting in increased accessibility of N-terminal galactose used by AAV9. Right: HeLa cells were treated with the indicated concentration of Neuraminidase 1hr prior to transduction with 1E4 vg/cell AAV9 or VCAP-102 containing a luciferase transgene. Luciferase assay was then performed 24hr post-transduction. Data indicate mean ± SD (n=3).

**Extended Data Fig. 8.**
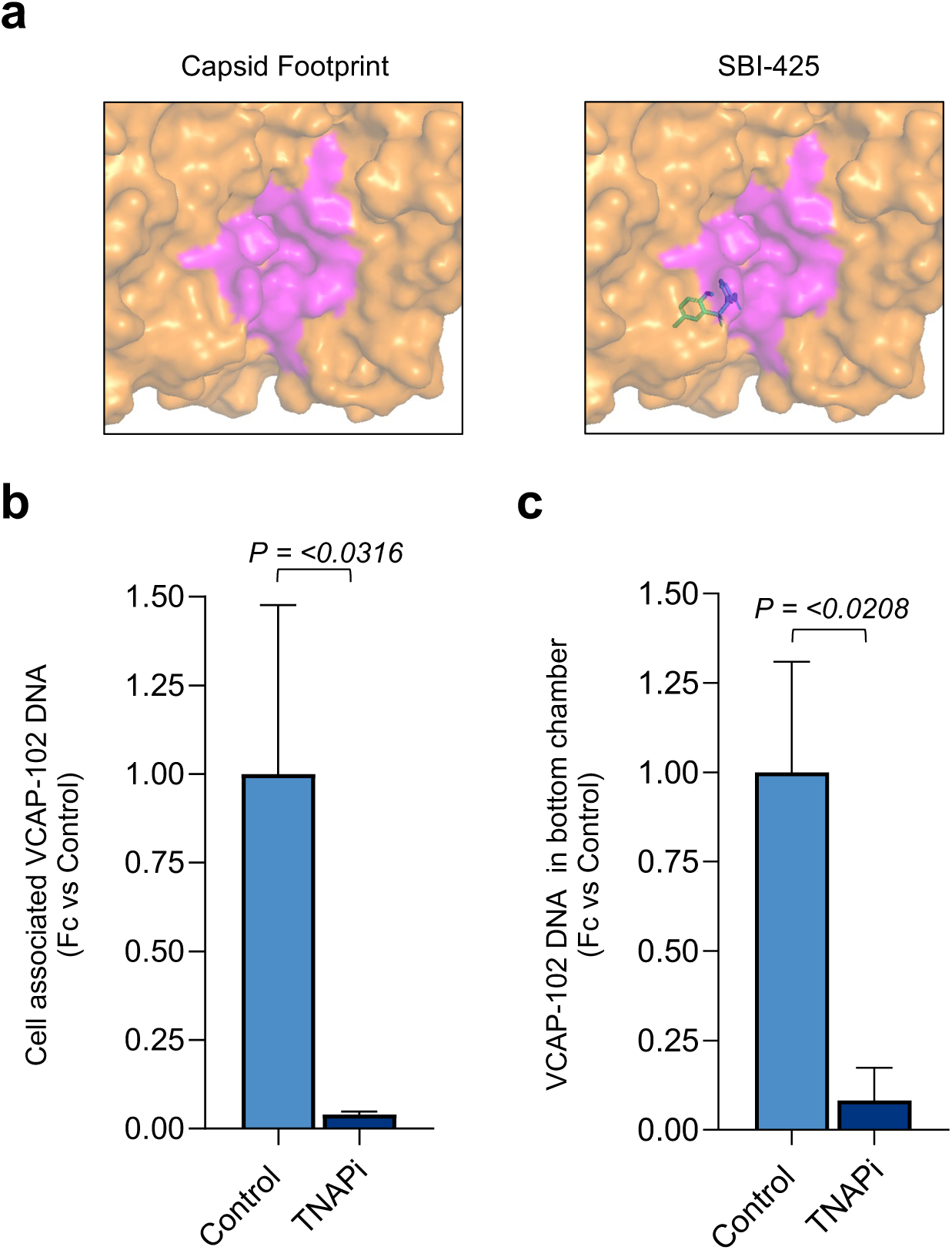
ALPL inhibitor disrupts capsid-receptor interaction and transcytosis. a, Top-down view of ALPL (orange) showing *in silico* prediction of VCAP-102 capsid footprint (magenta) and SBI-425 binding site (right panel), modeled with Diffdock. **b,** Quantification of VCAP-102 bound to MDCK-ALPL cells in the presence of 100 µM ALPL small molecule inhibitor (TNAPi). AAV genomes were quantified by Real-time RT-PCR, and data was normalized to untreated controls. **c,** Inhibition of VCAP-102 transcytosis by ALPL inhibitor. Transwell assay was performed with MDCK-ALPL in the presence of 100 µM of TNAPi. Viral genomes in the bottom chamber were quantified by qPCR. Data indicate mean ± SD normalized to untreated cells (n=3).

**Supplementary Table 1.** VCAP-102 dose-response quantification in mice. Automated quantification of cells in the mouse cortex positive for histone-HA as measured by co-localization of chromogenic HA staining and hematoxylin or by co-staining with cell-type specific markers and imaging through confocal microscopy. Note that each value in the table represents the average of three individual measurements within each mouse cortex.

|  |  |  | Chromogenic |  | Fluorescence |  |  |  |  |  |
| --- | --- | --- | --- | --- | --- | --- | --- | --- | --- | --- |
| Capsid: Transgene | Dosage (VG/kg) | Tissue | HA Positive Cells/mm<sup>2</sup> | % HA Positive Cells | HA Positive Cells/mm<sup>2</sup> | % HA Positive Cells | NeuN Positive Cells that are also HA Positive/mm<sup>2</sup> | % NeuN Positive Cells that are also HA Positive | Sox9 Positive Cells that are also HA Positive/mm<sup>2</sup> | % Sox9 Positive Cells that are also HA Positive |
| VCAP-102:CAG-ssH3-HA | 1.0E+14 | Cortex mouse 1 | 611.3 | 35.9 | 461.6 | 27.8 | 186.3 | 15.4 | 20.0 | 11.9 |
|  |  | Cortex mouse 2 | 781.3 | 40.8 | 549.0 | 31.2 | 151.0 | 13.3 | 38.9 | 16.9 |
|  |  | Cortex mouse 3 | 644.8 | 36.3 | 477.1 | 26.8 | 126.8 | 11.8 | 19.7 | 9.5 |
|  |  | **Average** | **679.1** | **37.7** | **495.9** | **28.6** | **154.7** | **13.5** | **26.2** | **12.8** |
|  | 3.2E+13 | Cortex mouse 1 | 367.6 | 22.2 | 266.7 | 16.7 | 144.5 | 10.7 | 14.6 | 17.9 |
|  |  | Cortex mouse 2 | 500.5 | 28.8 | 426.3 | 20.0 | 143.5 | 9.5 | 28.5 | 15.9 |
|  |  | Cortex mouse 3 | 374.1 | 27.8 | 338.9 | 19.6 | 92.2 | 7.6 | 16.1 | 9.9 |
|  |  | **Average** | **414.0** | **26.3** | **344.0** | **18.8** | **126.8** | **9.2** | **19.8** | **14.6** |
|  | 1.0E+13 | Cortex mouse 1 | 254.0 | 19.0 | 271.3 | 15.1 | 107.9 | 7.4 | 7.6 | 5.3 |
|  |  | Cortex mouse 2 | 363.2 | 23.3 | 312.6 | 18.0 | 66.4 | 5.5 | 15.5 | 10.0 |
|  |  | Cortex mouse 3 | 215.2 | 14.9 | 228.4 | 13.7 | 45.4 | 3.7 | 12.8 | 8.7 |
|  |  | **Average** | **277.5** | **19.1** | **270.7** | **15.6** | **73.2** | **5.5** | **12.0** | **8.0** |
|  | 3.2E+12 | Cortex mouse 1 | 131.0 | 8.3 | 151.3 | 8.4 | 49.0 | 3.3 | 10.9 | 7.1 |
|  |  | Cortex mouse 2 | 162.7 | 9.0 | 173.4 | 8.1 | 61.0 | 3.6 | 5.8 | 3.5 |
|  |  | Cortex mouse 3 | 161.8 | 8.1 | 206.0 | 8.0 | 67.6 | 3.9 | 8.8 | 2.7 |
|  |  | **Average** | **151.8** | **8.5** | **176.9** | **8.2** | **59.2** | **3.6** | **8.5** | **4.4** |
|  | 1.0E+12 | Cortex mouse 1 | 78.7 | 4.7 | 71.1 | 4.5 | 15.4 | 1.2 | 7.6 | 4.1 |
|  |  | Cortex mouse 2 | 49.4 | 2.6 | 43.9 | 2.5 | 2.3 | 0.2 | 1.7 | 0.8 |
|  |  | Cortex mouse 3 | 55.0 | 3.5 | 58.6 | 3.6 | 4.2 | 0.4 | 3.9 | 2.1 |
|  |  | **Average** | **61.0** | **3.6** | **57.9** | **3.5** | **7.3** | **0.6** | **4.4** | **2.3** |
| Vehicle | N/A | Cortex mouse 1 | 0.2 | 0.0 | 0.0 | 0.0 | 0.0 | 0.0 | 0.0 | 0.0 |
|  |  | Cortex mouse 2 | 0.2 | 0.0 | 0.0 | 0.0 | 0.0 | 0.0 | 0.0 | 0.0 |
|  |  | Cortex mouse 3 | 0.1 | 0.0 | 0.0 | 0.0 | 0.0 | 0.0 | 0.0 | 0.0 |
|  |  | **Average** | **0.2** | **0.0** | **0.0** | **0.0** | **0.0** | **0.0** | **0.0** | **0.0** |

**Supplementary Table 2.**
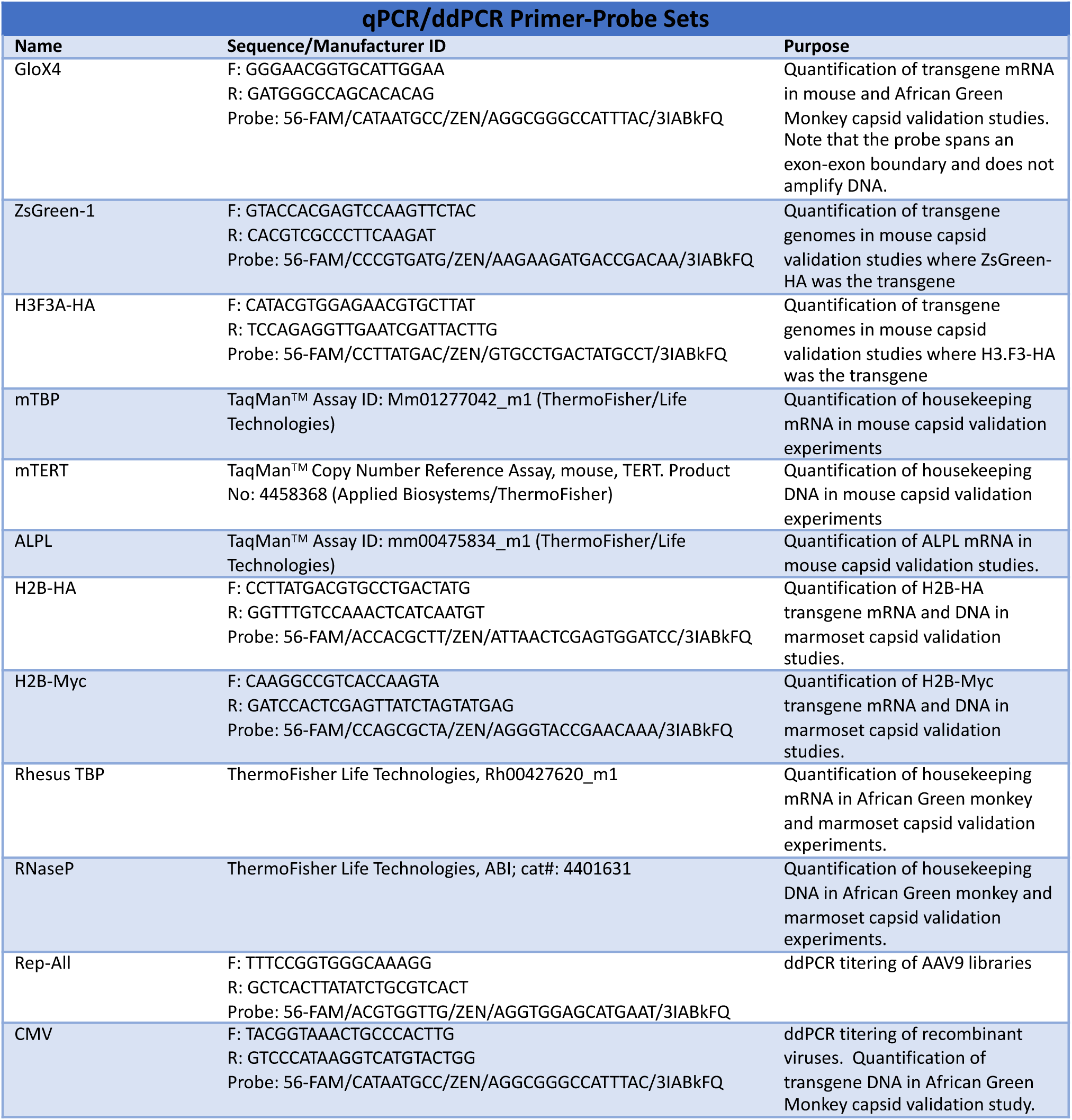
Key Reagents.

